# Anaerobic oxidation of methane in sediments of a nitrate-rich, oligo-mesotrophic boreal lake

**DOI:** 10.1101/2021.02.12.426818

**Authors:** Antti J Rissanen, Tom Jilbert, Asko Simojoki, Rahul Mangayil, Sanni L Aalto, Sari Peura, Helena Jäntti

**Affiliations:** Faculty of Engineering and Natural Sciences, Tampere University, Korkeakoulunkatu 6, FI-33720, Tampere, Finland; Ecosystems and Environment Research Program, Faculty of Biological and Environmental Sciences, University of Helsinki, P.O. Box 65, FI-00014, Helsinki, Finland; Department of Agricultural Sciences (Environmental Soil Science), Faculty of Agriculture and Forestry, University of Helsinki, P.O. Box 56, FI-00014, Helsinki, Finland; Department of Environmental and Biological Sciences, University of Eastern Finland, Yliopistonranta 1 E, FI-70210, Kuopio, Finland; Department of Biological and Environmental Sciences, University of Jyväskylä, Survontie 9 C, FI-40014, Jyväskylä, Finland; Department of Forest Mycology and Plant Pathology, Science for Life Laboratory, Swedish University of Agricultural Sciences, Almas allé 5, SE-75651, Uppsala, Sweden

**Author notes:** Corresponding author: Antti J Rissanen, Faculty of Engineering and Natural Sciences, Tampere University, Korkeakoulunkatu 6, FI-33720, Tampere, Finland. Technical University of Denmark, DTU Aqua, Section for Aquaculture, The North Sea Research Centre, P.O. Box 101, DK-9850, Hirtshals, Denmark.

**Keywords:** anaerobic oxidation of methane, nitrate, lake, methanotroph, shotgun metagenomics, 16S rRNA gene

## Abstract

The identity of electron acceptors in promoting anaerobic oxidation of methane (AOM) in the sediments of boreal lakes is currently unknown. Here, we studied the AOM rate of sediment slurries collected from three profundal stations of a nitrate-rich, oligo-mesotrophic, boreal lake (Lake Pääjärvi, Finland), under varying nitrate concentrations using ^13^C-labelling. Furthermore, vertical profiles of the sediment and porewater geochemistry, and the microbial communities (16S rRNA gene and shotgun metagenomic sequencing) were analyzed. Despite geochemical data indicating that simultaneous consumption of nitrate and methane took place at the sediment layers chosen for incubations, AOM rate was not enhanced by nitrate amendments at either of the stations. AOM rate was much higher at the shallow Station 1 (0.9-6.8 nmol C cm^-3^ d^-1^) with high contents of labile phytoplankton-derived organic matter, than at the deeper stations, Station 2 (0-0.3 nmol C cm^-3^ d^-1^) and 3 (0-0.2 nmol C cm^-3^ d^-1^). Accordingly, a higher relative abundance of methanotrophic archaea (*Candidatus* Methanoperedens) and bacteria (*Methylococcales*) were observed in the layers chosen for incubations at Station 1 than at the other stations. Besides nitrate, the geochemical profiles indicated that AOM was potentially coupled with iron or sulfate reduction at all stations. Furthermore, putative nitrite-reducing methanotrophs (*Ca*. Methylomirabilis) were the most abundant methanotrophs above the incubation layer at Station 2 and 3, which suggests that nitrite reduction also plays a role in driving AOM in the study lake. We conclude that AOM is not uniquely coupled to nitrate reduction in sediments of nitrate-rich, oligo-mesotrophic, boreal lakes.

## Introduction

The concentration of atmospheric methane (CH_4_), a critical greenhouse gas, has increased substantially since industrialization, with current total emissions 550 – 600 Tg/year (top-down estimates by Saunois et al. 2020). Roughly 40 % of these emissions stem from natural sources (Saunois et al. 2020), mostly produced by archaea through methanogenesis. Lakes are important natural emitters of CH_4_ (Bastviken et al. 2011). The numerous lakes and ponds in the northern boreal zone with annual emissions of ∼16.5 Tg are especially significant contributors to the global CH_4_ budget (Wik et al. 2016). Thus, knowledge on the factors affecting CH_4_ emissions from northern lakes is essential for accurate estimates and modelling of global CH_4_ budget.

Besides methanogenesis, microbial oxidation of CH_4_ taking place in sediments and water columns is an important controller of the CH_4_ emissions from lake ecosystems (Deutzmann et al. 2014). A recent study has reported that the anaerobic oxidation of methane (AOM) consumes large proportions (12 – 87 %) of gross CH_4_ production in sediments of some boreal lakes (Martinez-Cruz et al. 2018). Nevertheless, contrasting observations (i.e. negligible role of AOM in reducing CH_4_ emissions) from other boreal lakes have been also reported (Rissanen et al. 2017; Kallistova et al. 2018). This discrepancy can be due to varying availabilities of AOM-driving electron acceptors (EAs, i.e. NO_3_^-^, NO_2_^-^, Fe^3+^, Mn^4+^, SO_4_^2-^ and organic EAs) between lakes. However, the identity of EAs driving AOM in lake sediments is understudied and such data is especially scarce from boreal lakes (Rissanen et al. 2017; Kallistova et al. 2018; Martinez-Cruz et al. 2018). The quality of EAs driving AOM in natural ecosystems is usually studied in experimental incubations, where AOM rates are measured at increased concentration levels of EAs (Beal et al. 2009; Rissanen et al. 2017). In the few previous studies of boreal lake sediments, additions of Fe^3+^, Mn^4+^, SO_4_^2-^ or NO_3_^-^ did not enhance AOM (Rissanen et al. 2017; Kallistova et al. 2018). This was despite the fact that the sediments harbored typical freshwater AOM organisms, i.e. archaea in genus *Ca*. Methanoperedens sp. (also known as ANME-2D archaea), that potentially use a wide variety of EAs (e.g. NO_3_^-^, Fe^3+^, Mn^4+^ and SO_4_^2-^), and bacteria of genus *Ca*. Methylomirabilis sp. (within phylum *NC10*), that use NO_2_^-^ (Ettwig et al. 2010, 2016; Haroon et al. 2013; Weber et al. 2017; Rissanen et al. 2017). Previously, NO_3_^-^/NO_2_^-^ was shown to drive AOM in sediments of a eutrophic NO_3_^-^ -rich lake and a pond in temperate area (Norði and Thamdrup 2014; Deutzmann et al. 2014). However, such data do not exist from NO_3_^-^ -rich boreal lakes.

To shed light on the identity of EAs driving AOM in boreal lakes, we studied the effect of NO_3_^-^ additions on AOM in sediments of a NO_3_^-^ -rich boreal lake, Lake Pääjärvi (Southern Finland). This was complemented with analyses of vertical profiles of sediment and porewater geochemistry, and the associated microbial communities were studied using 16S rRNA gene sequencing and shotgun metagenomics. We hypothesized that due to the high availability of NO_3_^-^, AOM is linked to NO_3_^-^ reduction, and, thus, NO_3_^-^ addition enhances AOM in the sediments of the studied lake.

## Materials and methods

### Study lake

Lake Pääjärvi is a NO_3_^-^ -rich, oligo-mesotrophic lake in southern Finland (61.04N, 25.08E; A = 13.4 km^2^, max. depth = 87 m, mean residence time = 3.3 years). The water column circulates twice per year and is always well oxygenated (Salonen 1981). The large catchment area of Lake Pääjärvi (244 km^2^) is dominated by forests and agriculture. Since the 1970s, the nutrient concentrations of the water have increased (Arvola et al. 1996).

### Porewater and sediment sampling

Vertical profiles of the porewater and sediment samples were collected from three stations in Lake Pääjärvi on 9^th^ August 2017 using a hand-held HTH/Kajak corer with plexiglass tubes (Table 1). The stations followed a water depth gradient, Station 1 being shallowest and Station 3 deepest (Table 1). The core tubes were pre-drilled with two vertical series of 4 mm holes (each at 2 cm resolution), then taped, in preparation for porewater sampling with Rhizons^™^ (Rhizosphere research products, Wageningen, Netherlands). Rhizon sampling automatically filters the porewaters at 0.15 µm into attached 10 ml syringes under vacuum. One vertical series of samples was taken for analysis of dissolved CH_4_ and dissolved inorganic carbon (DIC). The syringes were pre-filled with 1 ml of 0.1M HNO_3_ to immediately convert all DIC to CO_2_. A known volume of N_2_ gas headspace was injected into the syringes after sampling, and the samples were shaken to equilibrate the dissolved gases with the headspace. The subsamples of the headspace were then extracted into 3 mL Exetainers^™^ (Labco Limited, Lampeter, UK) and stored at room temperature until analysis (for full details see Jilbert et al. 2020). The second vertical series was taken for analyses of S, Fe, and P [by inductively coupled plasma atomic emission spectroscopy (ICP-OES)], NO_3_^-^, and organic fermentation products. Subsamples for analyses of NO_3_^-^ and fermentation products were stored frozen at −20 °C until analysis. The subsamples for ICP-OES were acidified with 1 M HNO_3_ and stored at room temperature. After porewater sampling, the sediment cores were sliced into plastic bags at a resolution of 1 cm or 2 cm at 0-10 cm or >10 cm depth below the surface of sediment, respectively. The subsamples of 400 - 500 µl wet sediment were collected from the depth of 0-10 cm at each station and stored frozen at −20 °C for DNA-based molecular microbiological analyses. The remaining wet sediment samples were stored frozen at −20 °C under N_2_ until further processing.

**Table 1.** Station coordinates, depth, temperature and O_2_ concentration measured approximately 50 cm above the sediment surface and NO_3_ ^-^ concentrations measured 5 cm above the sediment. Loss on ignition (LOI) was measured from the top 0.5 cm of the sediment

| Station | N | E | Depth<br>(m) | Temp<br>(°C) | O<sub>2</sub><br>(μmol L<sup>-1</sup>) | NO<sub>3</sub><sup>-</sup><br>(μmol L<sup>-1</sup>) | LOI<br>(%) |
| --- | --- | --- | --- | --- | --- | --- | --- |
| 1 | 61.03.046 | 25.03.606 | 14 | 9.3 | 254 | 60.0 | 14.6 |
| 2 | 61.03.192 | 25.05.449 | 22 | 6.4 | 314 | 62.0 | 13.4 |
| 3 | 61.03.487 | 25.06.163 | 52 | 5.4 | 303 | 45.5 | 14.2 |

### Porewater and sediment bulk geochemical analysis

According to our standard protocols, all the Rhizon samples were sequentially washed with 0.01 M HNO_3_ and MilliQ water before field sampling. Later, it was noticed that the washing procedure caused background contamination in the porewater NO_3_^-^ measurements. The magnitude and variability of this contamination was tested by sampling a solution of MilliQ water through the Rhizon samplers in plastic syringes and determining the NO_3_^-^ concentration of the extracted sample. The same procedure was repeated using both factory-clean (9 replicates) and 0.01M HNO_3_-washed Rhizons (10 replicates). The difference between the mean NO_3_^-^ values, determined from the factory-clean and washed Rhizon samples (30.0 µmol L^-1^), was subtracted from all the porewater NO_3_^-^ data of this study. For objectivity, we report here the corrected values, as well as the error margins representing the full range of observed contamination (min – max, 16.8 µmol L^-1^ - 73.9 µmol L^-1^). The porewater nitrate concentrations were determined according to Miranda et al. (2001), while the samples from the HNO_3_ – contamination test were analyzed by QuikChem®8000 flow injection analyzer (Lachat Instruments, Hach Co., Loveland, CO, USA) according to Wood et al. (1967).

The S, Fe and P concentrations in the porewater samples were determined by ICP-OES (Thermo iCAP 6000, Thermo Fisher Scientific, Waltham, MA, USA). In this system, S is expected to be dominated by sulfate (SO_4_^2-^), Fe by Fe^2+^, and P by orthophosphate (H_2_PO_4_^-^), although in each case, other forms are possible. The porewater CH_4_ and DIC concentrations were determined by gas chromatography (GC) as described in Jilbert et al. (2020). Briefly, the sample vials (Exetainers) were pressurized with helium (He) to 2.0 bar before loading into the GC (Agilent technologies 7890B GC system, Agilent Technologies, Santa Clara, Ca, USA). CH_4_ was determined by flame-ionization detector (FID) and CO_2_ by thermal conductivity detector (TCD). The instrument simultaneously measures N_2_ and O_2_ + Ar (in TCD), from which a 100% sum can be calculated for estimation of CH_4_ and CO_2_ concentrations in the original sample, in ppm by volume. For a full description of the calculations, see Jilbert et al. (2020).

Sediment samples were freeze-dried, ground in an agate mortar and weighed into tin cups for carbon (C) and nitrogen (N) content determinations. In accordance with extensive previous studies on Finnish lake sediments (Pajunen 2000; Kortelainen et al. 2004), acidification was not applied prior to the determinations. High levels of organic acidity from Finnish river catchments (Kortelainen and Saukkonen 1995) maintain low annual mean pH values in most lakes and therefore there is negligible occurrence of carbonates in lake sediments. Hence our total C data is considered equivalent to organic C (C_org_). C and N contents were determined using elemental analyzer (LECO TruSpec Micro, LECO Corp., St. Joseph, MI, USA).

Organic fermentation products in porewaters were analyzed using high performance liquid chromatography (HPLC) equipped with Shodex SUGAR column (300 mm × 8 mm, Phenomenex, Torrance, CA, USA), autosampler (SIL-20AC HT, Shimadzu, Kioto, Japan), refractive index detector (RID-10A, Shimadzu), and 0.01 M H_2_SO_4_ as the mobile phase. The HPLC samples were prepared as described in Salmela et al. (2018). The identification and quantification of the liquid metabolites (lactate, acetate and propionate) were conducted using external standards.

### Incubations for the AOM measurements

The sediment for the incubation experiments was collected from the same three stations, one week after the collection of porewater and sediment samples (15^th^ Aug 2017), using plexiglass tubes (diameter = 7 cm) connected to a gravity corer (Table 1). The initial porewater profile data were used to select the optimum depth horizons for sampling sediments for the incubations. At each station, a 3 cm thick sediment slice was cut immediately below the layer at which the porewater Fe concentrations started to increase downwards, indicating the onset of anaerobic remineralization processes and therefore a suitable depth interval for the incubation experiments. The upper limits to the incubation slices are as follows: −1 cm at Station 1 (thus, layer 1 – 4 cm from surface); −3 cm at Station 2 (layer 3 – 6 cm) and −3 cm at Station 3 (layer 3 – 6 cm).

The sediment from each station was mixed in 1:1 ratio with filtered (0.6 µm glass fiber filter) water collected 50 cm above the sediment. The sediment slurry was prepared in a 1 L bottle and purged with N_2_ for 10 min. Thereafter, the bottle was quickly capped with a septum and the headspace was purged again with N_2_ for 5 min. To ensure anoxic conditions prior to the experiments, the bottle containing the sediment slurry was placed on a shaker table overnight at 5°C and 120 rpm to facilitate the removal of any residual O_2_. Thereafter, the set-up of incubations was conducted in a glove bag with N_2_ atmosphere. The sediment slurry was transferred into 12 ml glass vials (Exetainers) (2 ml of slurry per vial), after which the vials were filled with filtered (0.6 µm), anoxic, N_2_-purged bottom water. Thereafter, sterile, anoxic, N_2_-purged stock solution of KNO_3_ was added to the vials in concentrations of 0, 10, 50, 100, 200 and 500 µmol (N) L^-1^. There were 57 vials altogether: i.e. three vials per each NO_3_^-^ concentration at each station (3 stations x 6 concentrations ⨯ 3 vials = 54) and, furthermore, one vial at each station represented a non-incubated control sample (without KNO_3_) of the slurry at the beginning of the experiment (= 3). The vials were capped with septum, followed by the addition of 100 µl of gaseous ^13^C-labeled CH_4_ (^13^C = 99.9%, Cambridge Isotope Laboratories Inc., Tewksbury, MA, USA) into two vials per each NO_3_^-^ concentration at each station. One vial per each NO_3_^-^ concentration at each station was not amended with ^13^C-CH_4_ and hence was included as a non-labeled incubation control. The vials were taken out of the glove bag, shaken for 3 minutes to aid the dissolution of the added methane and incubated on a shaker (120 rpm) at 5 °C in the dark for 21.7, 20.75 and 20.25 hours for Station 1, 2 and 3, respectively.

After incubations, 3 ml of slurry per sample were injected into pre-evacuated He-filled 12 ml vials (Exetainers) (over-pressure was released from the He-filled vials before sampling), which contained 300 μL of H_3_PO_4_ (85%) to ensure the transformation of DIC into CO_2_. The samples were then stored inverted at +4 °C until the analysis.

### Stable isotopic and concentration analyses of DIC

Stable isotopic and concentration analyses of DIC were performed using an Isoprime100 isotope ratio mass spectrometer (IRMS) (Elementar UK Ltd., Cheadle, UK) coupled to an Isoprime TraceGas pre-concentration unit. The isotopic data was calculated as the fractional abundance of ^13^C, i.e. ^13^F, in DIC, using equation 1.

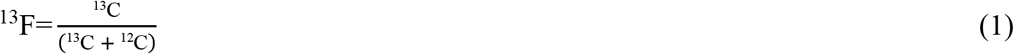

### Calculations

A simple two-component mixing model was applied for a first-order quantification of the relative contributions of terrestrial plant-derived and autochtonous phytoplankton-derived organic matter to the total sedimentary organic matter (% of total organic C). The calculation uses the molar N:C ratio of organic matter, and end-member values, N/C_EM_, based on the studies of Goñi et al. (2003) and Jilbert et al. (2018):

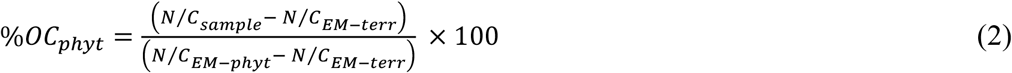

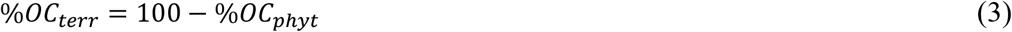

where N/C_EM-terr_ = 0.04, and N/C_EM-phyt_ = 0.13. The calculation assumes that terrestrial plant matter and phytoplankton are the only sources of organic material present in the samples, the N/C values are spatially and temporally fixed at the end-member values, and the values do not alter significantly during sedimentation and burial of organic matter.

AOM rates were determined from the production rate of excess ^13^C-labeled DIC,

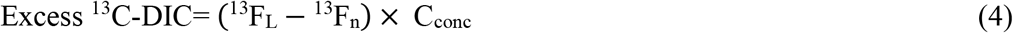

where C_conc_ is the concentration of DIC, and ^13^F_L_ is the fractional abundance of ^13^C in ^13^C-labeled samples, while ^13^F_n_ denotes that in non-labeled samples (see equation 1). Assuming that the excess ^13^C-DIC was zero at the beginning, we used a single time point measurement, i.e. excess ^13^C-DIC after the incubations, to determine the AOM rate, expressed as a rate per dry weight (DW) or per volume of sediment per hour (nmol C g^-1^_DW_ h^-1^ or nmol C cm^-3^ h^-1^). The added amount of CH_4_ (i.e. 100 µl, see above) corresponds to a concentration of approximately 320 µmol L^-1^, which is on average 8 – 14 times higher than the natural CH_4_ concentration in the depth layers chosen for the incubations (see Results and Discussion). Furthermore, considering that the slurry preparations led to only 1 mL of the original sediment to be added to each vial (i.e. 2 mL of 1:1 water:sediment – slurry added to each 12 mL vial, which was then filled with water, see above), the added ^13^C-labeled CH_4_ was estimated to account of almost all (> 99 %) of CH_4_ in the incubation vial. Thus, the measured production rates of excess ^13^C-DIC represent AOM rates. The CH_4_ oxidation in this study was determined solely based on the transfer of ^13^C from CH_4_ to DIC, and hence, the proportion of CH_4_-C bound to the biomass or extracellular organic metabolites was not taken into account.

### DNA extraction

DNA was extracted from the frozen sediment samples using DNeasy PowerSoil Kit (Qiagen, Hilden, Germany). DNA concentration was measured using a Qubit 2.0 Fluorometer and a dsDNA HS Assay Kit (Thermo Fisher Scientific, Waltham, MA, USA).

### 16S rRNA gene amplicon sequencing

PCR and 16S rRNA gene amplicon sequencing was performed by The Foundation for the Promotion of Health and Biomedical Research of Valencia Region (FISABIO, Valencia, Spain). In the PCR reactions, the V4 region of the bacterial and archaeal 16S rRNA genes were simultaneously targeted using primer pair 515FB (5’-GTGYCAGCMGCCGCGGTAA-3’)/806FB (5’-GGACTACNVGGGTWTCTAAT-3’) (Apprill et al. 2015; Parada et al. 2016). PCR, library preparation and sequencing (Illumina MiSeq, Illumina, San Diego, CA, USA) was performed as previously described (Myllykangas et al. 2020), except that, in PCR reactions, approximately 15 ng of DNA were used.

### Shotgun metagenomic sequencing

The shotgun metagenomic sequencing was performed by Novogene Co., Ltd (Hong Kong). The libraries were prepared from ∼100 ng of DNA per sample using the NEBNext® Ultra™ DNA Library Prep Kit for Illumina® (New England Biolabs, Ipswich, MA, USA) according to the manufacturer’s protocol. The sequencing was done on Illumina HiSeq X Ten System (Illumina) with paired-end mode and 150 bp read length producing from ∼9 to ∼12 Gb of raw data per sample.

### Bioinformatic analyses of 16S rRNA gene data

The quality assessment of the raw sequence reads, merging of paired end reads, alignment, chimera removal, preclustering, taxonomic classification (and removal of chloroplast, mitochondria, and eukaryote sequences) and division of 16S rRNA gene sequences into operational taxonomic units (OTUs) at 97 % similarity level was conducted as previously described (Rissanen et al. 2020). Singleton OTUs (OTUs with only 1 sequence) were removed, and the data were then normalized by subsampling to the same size, 77340 sequences. One sample, representing the layer 8-9 cm depth at Station 3, was discarded from the analyses since it had only ∼10 000 sequences.

### Bioinformatic analysis of shotgun metagenomic data

The raw sequences were analyzed using The European Bioinformatics Institute (EBI) Metagenomics pipeline (version 4.1) for trimming and taxonomic analysis of the reads containing 16S rRNA gene sequences against Silva 132 database (Mitchell et al. 2017). Furthermore, Kaiju web server (http://kaiju.binf.ku.dk/server) was used for taxonomic classification of the raw reads against National Center for Biotechnology Information’s (NCBI) nr – database (Menzel et al. 2016).

For metagenomic binning analysis, the raw data was trimmed using Trimmomatic (version 0.39) (Bolger et al. 2014). The trimmed data was assembled using Megahit (version 1.2.8) (Li et al. 2015). The quality-controlled reads were mapped to the assembly using BBmap (version 38.73) (https://sourceforge.net/projects/bbmap/). The mapping results were used to bin the contigs using Metabat2 (Kang et al. 2015, 2019). The genes of the obtained metagenomic bins were predicted and annotated using Prokka (version 1.14.5) (Seemann 2014). The bins were also taxonomically annotated using CAT/BAT – tool and NCBI’s nr – database and taxonomy (von Meijenfeldt et al. 2019). Prokaryotic completeness and redundancy were computed using CheckM (version v1.1.2) (Parks et al. 2015). Taxonomic annotations were specifically screened for bins affiliated to known aerobic (gamma- and alphaproteobacterial MOBs) and anaerobic methanotrophs (ANME archaea and *Ca*. Methylomirabilis sp. bacteria). Furthermore, functional annotations were specifically screened for bins coding for particulate and soluble methane mono-oxygenases, key enzymes in aerobic methanotrophy and in *Ca*. Methylomirabilis sp. – mediated AOM, and for methyl coenzyme M reductase, the key enzyme in methanogenesis and in ANME archaea - mediated AOM.

### Statistical analyses

The variation between stations and between NO_3_^-^ concentrations was studied using a two-way analysis of variance (2-ANOVA) with station (3 levels) and NO_3_^-^ concentration (6 levels) as fixed factors followed by pair-wise post hoc tests using least significant difference (LSD) technique with Hochberg-Bonferroni corrected α-values. The relationship between the varying NO_3_^-^ concentration and AOM was also studied specifically at each station using Spearman correlation analysis.

In addition, the variation between stations in relative abundance of different methanotrophic, methanogenic and SO_4_^2-^ reducing taxa at the depth layers representing the AOM incubation depths was analyzed using randomized block analysis of variance (RB-ANOVA) with the depth layer (three 1 cm layers at each station) as the blocking factor. RB-ANOVA was followed by pair-wise post-hoc tests using LSD technique with Hochberg-Bonferroni-corrected α-values. Furthermore, the coherence in the depth patterns between ANME archaea and SO_4_^2-^ reducing bacteria was analyzed by Spearman correlation analysis. All the analyses were done using IBM SPSS Statistics 26 (IBM SPSS Statistics for Windows, Version 26.0, IBM Corp., Armonk, NY, USA).

### Sequence database accession

The sequences of the 16S rRNA gene amplicon dataset are deposited in NCBI’s Sequence Read Archive (SRA) under the accession number PRJNA627335. The sequences of the shotgun metagenomic dataset are deposited in European Nucleotide Archive (ENA) under the accession number PRJEB29513.

## Results and discussion

### Sediment chemical characteristics and the structure of the methanogenic community

All the stations showed a broadly similar diagenetic zonation based on the porewater chemistry (Figs. 1 and 2). The vertical profiles of porewater concentrations of NO_3_^-^, S (assumed to be mostly SO_4_^2-^), Fe (assumed to be mostly Fe^2+^ from reduction of solid-phase Fe oxides) and CH_4_ suggest a transition zone, wherein the upwards-diffusing CH_4_ is oxidized by anaerobic respiration linked to reduction of NO_3_^-^, SO_4_^2-^, and Fe oxides. This zone was slightly closer to the sediment surface at Station 1 than at the other stations (Figs. 1 and 2). Station 1 also had higher overall concentrations of CH_4_ and CO_2_ than Station 2 and 3, and was the only station where lactate was observed in the porewaters (Fig. 2). In addition, while acetate and propionate were observed at all stations, they were at slightly higher concentrations at Station 1 than at the other two stations (Fig. 2). These results suggest that there was lower redox-potential and higher degree of fermentation at Station 1, which very likely led to higher rates of production of fermentation products and methanogenesis. This may be due to higher lability of organic matter, being supported by our estimates on the contribution (%) of phytoplankton-derived organic carbon, which was slightly higher at Station 1 than at Station 2 and 3 (Fig. 2).

**Fig. 1.**
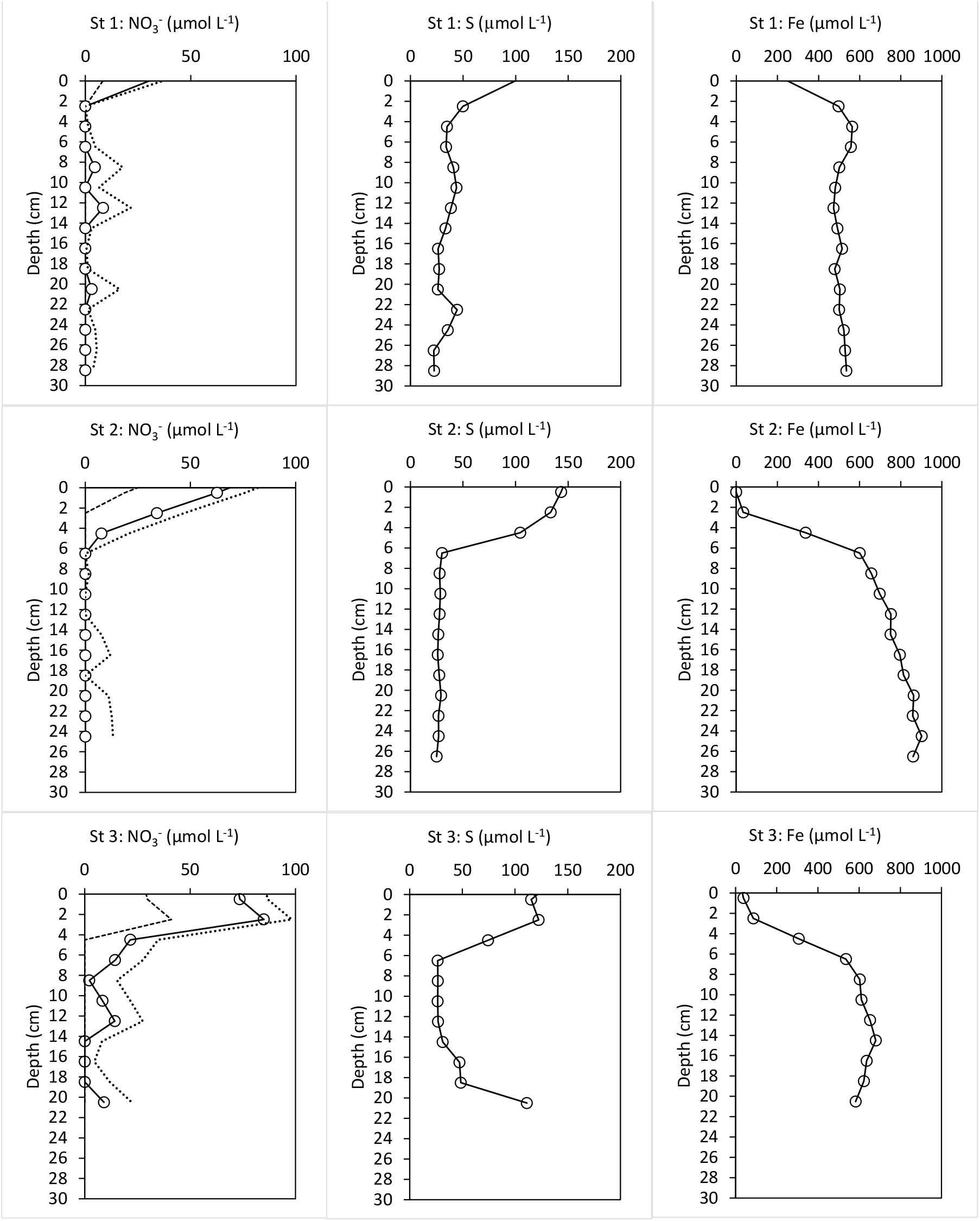
Sediment porewater profiles of dissolved NO_3_^-^ (left panels), S (middle panels) and Fe (right panels) at Stations 1 (upper panels), 2 (middle panels) and 3 (lower panels). The extent of the NO_3_^-^ contamination resulting from HNO_3_ washing of porewater samplers was tested and the NO_3_^-^ data is presented as values from which average (solid line with symbols), minimum and maximum contamination (dotted lines) was subtracted (see Materials and Methods). To allow comparison with the microbiological data, the profiles are specifically shown for the top 10 cm layer also in Fig. 3

**Fig. 2.**
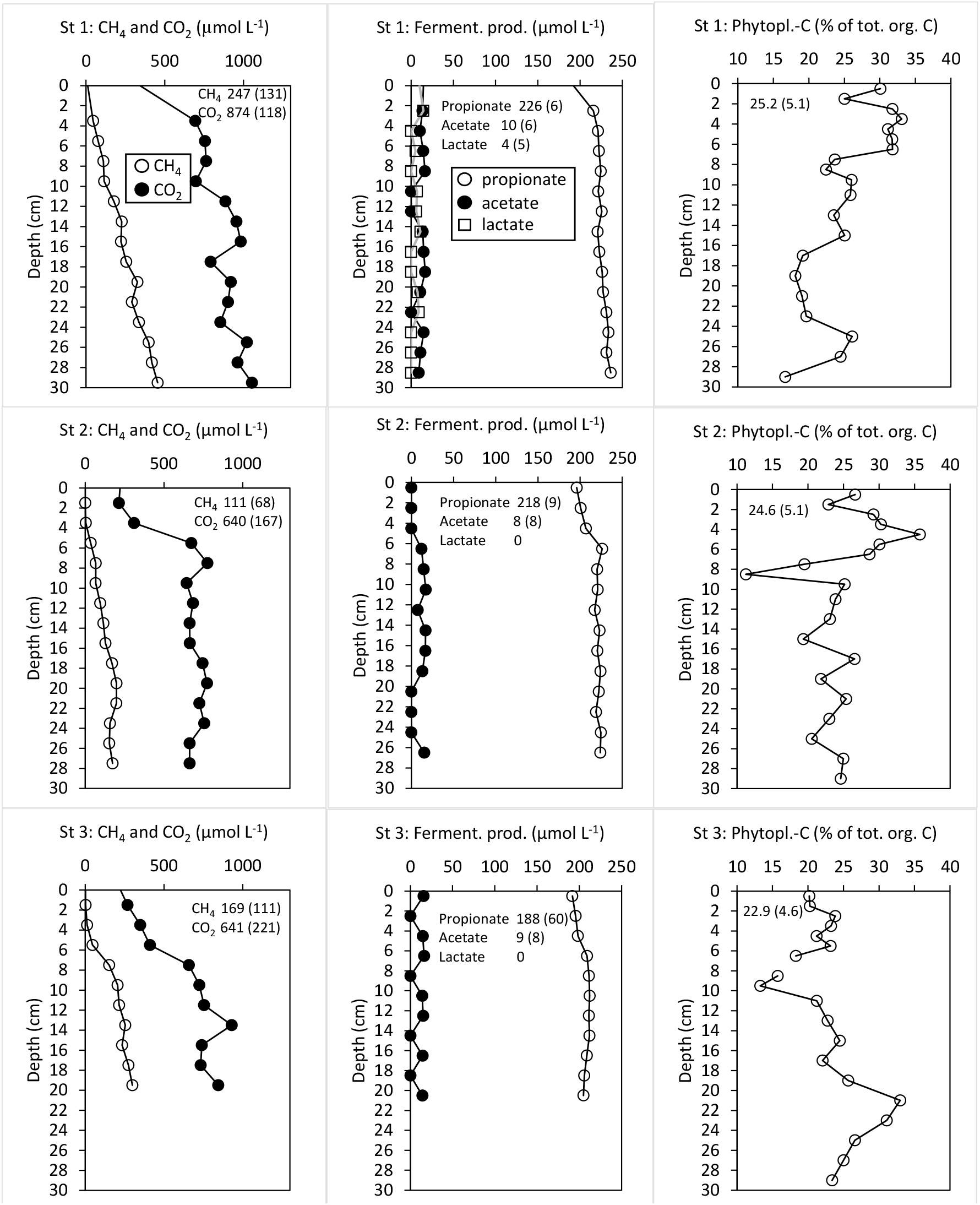
Profiles of CH_4_ and CO_2_ (left panels), organic fermentation products (middle panels) and estimated contribution of labile phytoplankton-derived organic carbon in total organic carbon in sediments (right panels) at Stations 1 (upper panels), 2 (middle panels) and 3 (lower panels). Average values (±SD) calculated across the profiles for each variable are shown within the figures. To allow comparison with the microbiological data, the profile of CH_4_ is specifically shown for the top 10 cm layer also in Fig. 3

In accordance with lower redox-potential and higher rates of methanogenesis at Station 1, the relative abundance of methanogens [shown as average % (min-max) of 16S rRNA gene amplicons] in the studied layer (topmost 10 cm) was generally higher there [0.5 % (0.01-1.1 %)] than at Station 2 [0.3 % (0.006-1.8 %)] and 3 [0.1 % (0.003-0.5 %)] (Fig. 3). The order *Methanomassiliicoccales*, whose cultivated members drive solely the methyl reducing methanogenic pathway (Borrel et al. 2013), dominated the methanogen communities at all stations (Fig. 3). Nevertheless, the hydrogenotrophic (i.e. H_2_-consuming) CO_2_ reducing methanogens of the order *Methanomicrobiales* (Nazaries et al. 2013) made also a significant proportion of methanogenic community at Station 1 but not at the other two stations (Fig. 3). The dominance of *Methanomassiliicoccales* is surprising, because methanogenesis in freshwater sediments is typically dominated by the acetoclastic (i.e. acetate – utilizing) and hydrogenotrophic CO_2_ reducing pathways (Conrad 1999). However, a recent metagenomic data suggested the presence of the acetoclastic pathway in genomes of uncultivated *Methanomassiliicoccales*, which could explain our findings (Speth and Orphan 2018). On the other hand, being only very recently described order with currently limited information on its functional diversity (Iino et al. 2013), *Methanomassiliicoccales* might also contain species without capability for methanogenesis. Unfortunately, we could not construct metagenome-assembled genomes (MAGs) of *Methanomassiliicoccales* or any other methanogens, probably because they had too low abundance to be detected by the metagenomic binning analysis (see Supplementary Results and Table S1). Hence, the potential functions of *Methanomassiliicoccales* in sediments of the study lake remain uncharacterized.

**Fig. 3.**
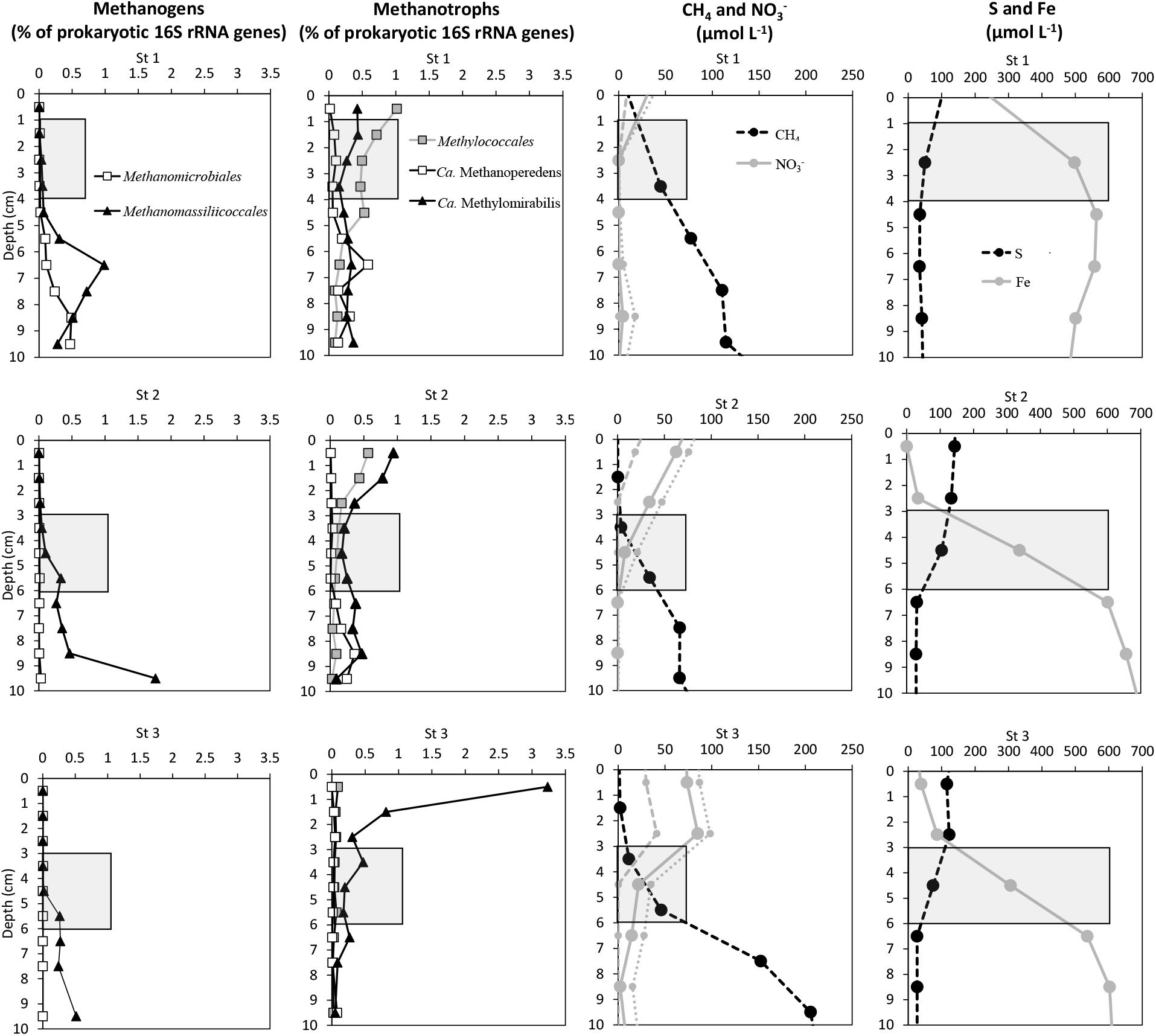
Vertical depth profiles of methanogens (left panels) and methanotrophs (second left panels) based on 16S rRNA gene amplicon sequencing, as well as CH_4_, NO_3_^-^ (second right panels), S and Fe (right panels) in the topmost 10 cm of sediments at Stations 1 (upper panels), 2 (middle panels) and 3 (lower panels). The layers chosen for AOM incubation analyses are highlighted with grey color. See Fig. 1 for explanation on the presentation of the NO_3_^-^ data

### AOM rates in the sediment

The effect of NO_3_^-^ addition on AOM rates in sediment slurries consisting of layer 1 – 4 cm (from surface) at Station 1 and layer 3 – 6 cm at Station 2 and 3 was analyzed by anaerobic incubations using ^13^C-labeled CH_4_. The complete pore water profiles show that the incubated layers of each core were suitably chosen at the transition between high NO_3_^-^ and S concentrations (above) and high Fe^2+^ and CH_4_ concentrations (below), indicating the zone of active reduction of NO_3_^-^, SO_4_^2-^ and Fe^3+^ and consumption of CH_4_ (Figs. 1-3).

In comparison to non-labelled control samples, incubation with amendments of ^13^C-labeled CH_4_ led to ^13^C-enrichment of DIC. The ^13^C-enrichment of DIC was much higher at Station 1 than at Station 2 and 3 (Fig. S1). In accordance, the AOM rates were significantly higher at Station 1 (min-max: 0.3 – 1.8 nmol C g^-1^_DW_ h^-1^) than at Station 2 (0 – 0.07 nmol C g^-1^_DW_ h^-1^) or 3 (0 – 0.06 nmol C g^-1^_DW_ h^-1^) (2-ANOVA, F_Station_ = 45, p < 0.001) (Fig. 4). The AOM rates (converted to volumetric rates, 0 – 6.8 nmol C cm^-3^ d^-1^) are within the lower end of AOM rates reported from lake sediments in general (0 – 180 nmol C cm^-3^ d^-1^) and from boreal lakes in particular (0 – 16.3 nmol C cm^-3^ d^-1^) (Deutzmann and Schink 2011; Rissanen et al. 2017; Martinez-Cruz et al. 2018). However, the AOM rates at Station 1 (0.9 – 6.8 nmol C cm^-3^ d^-1^) correspond well with the median AOM (3.5 nmol C cm^-3^ d^-1^) of the multiple lake studies reviewed by Martinez-Cruz et al. (2018).

**Fig. 4.**
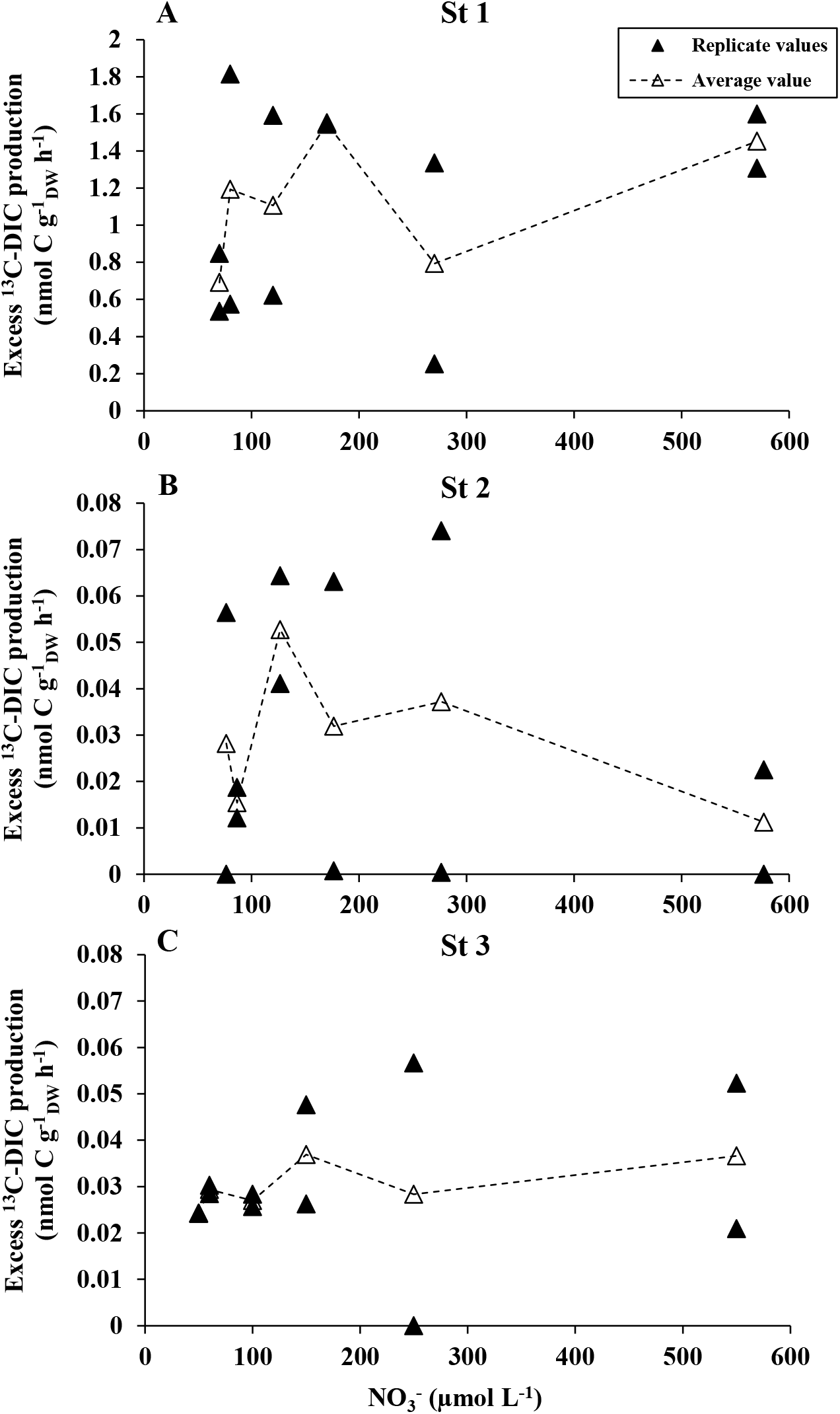
AOM rate, i.e. production of excess ^13^C-DIC, at different concentrations of NO_3_^-^ at **A** Station 1, **B** Station 2 and **C** Station 3. Values are represented as replicate and average values (n=2 at each concentration level of NO_3_^-^). The NO_3_^-^ concentration levels at the x-axis represent the sum of added NO_3_^-^ (i.e. 0, 10, 50, 100, 200 or 500 µmol L^-1^) and NO_3_^-^ already present in slurry at the beginning of incubations (i.e. 70, 77 and 50 µmol L^-1^ for stations 1, 2 and 3, respectively). No positive relationship between NO_3_^-^ and AOM was observed at the stations. Note different scale in y-axis in **A** compared to **B** and **C**

In agreement with both insignificant rank correlation (p > 0.05) and 2-ANOVA analyses (F_concentration_ = 0.7, p > 0.05), increasing concentration of NO_3_^-^ did not enhance AOM at any of the stations (Fig. 4). Hence, despite the evidence from the geochemical profiles, these results suggest that AOM was not directly coupled with NO_3_^-^ reduction in the sediments of the study lake. We acknowledge that the tested nitrate concentration levels (sum of added nitrate and nitrate already present in the slurry) were on average 7-57, 5-35 and 1-13 times higher than that naturally present in the incubation layers at Station 1, 2 and 3, respectively (Figs. 3 and 4). However, we find it unlikely that the observed independence between nitrate concentration and AOM would be caused by nitrate-saturation of AOM at high nitrate concentrations. This is because the fact that also the added ^13^C-CH_4_ concentrations were much higher, on average 9, 14 and 8 times higher, than natural CH_4_ concentrations at Station 1, 2 and 3, respectively. Hence, instead of NO_3_^-^, AOM appears to be coupled with reduction of other EAs, e.g. Fe^3+^ or SO_4_^2-^, as suggested by the geochemical profiles, or organic EAs (Valenzuela et al. 2017; Su et al. 2019). Furthermore, under SO_4_^2-^ -starved conditions, Fe^3+^ reduction can also indirectly support SO_4_^2-^ -mediated AOM through oxidation of reduced sulfur species to SO_4_^2-^ (Su et al. 2019).

### Methanotrophic community in the sediments

Aerobic MOBs, in order *Methylococcales*, as well as anaerobic methanotrophs, in genera *Ca*. Methanoperedens sp. (also known as ANME 2D archaea) and *Ca*. Methylomirabilis sp. (within bacterial phylum *NC10*), were the most abundant groups of methanotrophs present in the studied sediments, agreeing with a previous study on boreal lake sediments (Rissanen et al. 2017) (Figs. 3 and 5). Aerobic MOBs in the family *Methylacidiphilaceae* were rare, while alphaproteobacterial MOBs were not detected. As expected for aerobic bacteria, the relative abundance of *Methylococcales* methanotrophs was highest at the surface sediment layers and decreased towards deeper layers (Fig. 3). In contrast, *Ca*. Methanoperedens sp. had peaks in its relative abundance at deeper layers clearly below the sediment surface (Fig. 3). *Ca*. Methylomirabilis sp. peaked in its relative abundance both at the surface as well as at deeper sediment layers at all stations (Fig. 3).

**Fig. 5.**
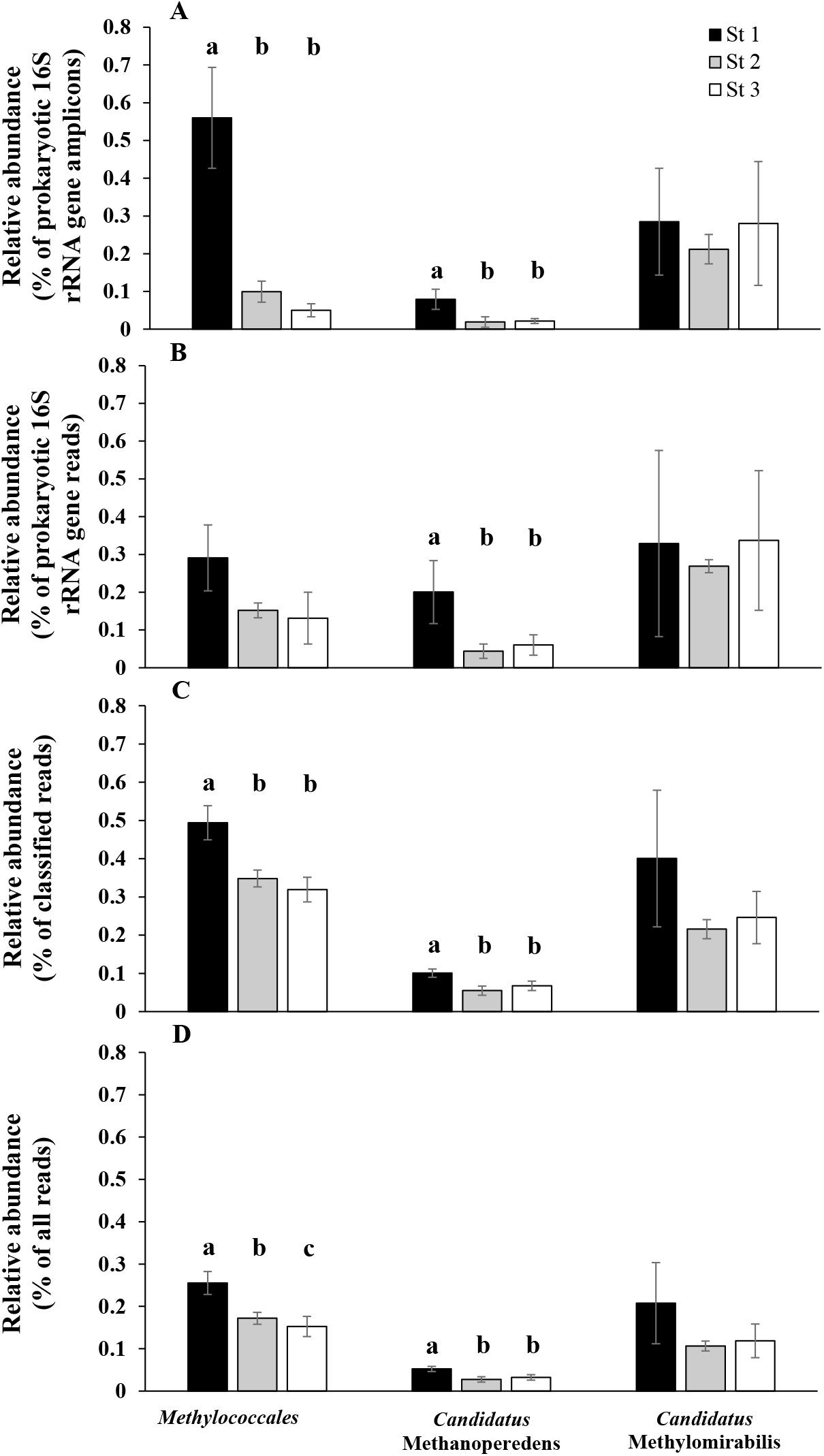
Relative abundance of different methanotrophic taxa in the depth layers representing the incubation layers based on **A** 16S rRNA gene amplicon sequencing, as well as **B**-**D** shotgun metagenomic sequencing, i.e **B** 16S rRNA gene reads, **C** KAIJU-classified reads expressed as relative to reads that were classified at least to domain-level and **D** KAIJU-classified reads expressed as relative to all reads. Data is represented as an average and standard deviation of three layer-specific samples. Different letters above bars indicate significant differences in RB-ANOVA and subsequent pairwise tests (p < 0.05)

Methanotroph communities in the depth layers representing the layers chosen for AOM incubation were specifically compared between stations. As the results of amplicon sequence analyses can be affected by PCR-based biases (e.g. primer bias), these comparisons were also made using PCR-free shotgun metagenomic techniques. *Ca*. Methanoperedens sp. and *Methylococcales* had higher relative abundances at Station 1 than at other stations (Fig. 5), which validates the higher AOM rates obtained from Station 1 (Fig. 4). In contrast, the relative abundance of *Ca*. Methylomirabilis did not vary between the stations (Fig. 5). We acknowledge that these comparisons are based on relative abundances and we lack analyses of absolute abundances (e.g. by quantitative PCR). However, we can assume that the relative abundance of a certain taxa, when expressed as the number of shotgun metagenomic reads relative to all sequenced reads, represents a proxy for the proportion of DNA of the taxa in the total extracted DNA (Fig. 5D). As concentration of DNA in the sediment (average ± SD of three layer specific samples), i.e. 1040 ± 290 ng cm^-3^, 1170 ± 470 ng cm^-3^ and 870 ± 100 ng cm^-3^ at Station 1, 2 and 3, respectively, did not differ between the stations (RB-ANOVA, F = 0.5 and p > 0.05), the variations in the relative abundances expressed as number of reads relative to all sequenced reads suggests that also the absolute abundance of *Ca*. Methanoperedens and *Methylococcales* was higher at Station 1 than at the other two stations (Fig. 5D).

In addition to using NO_3_^-^, *Ca*. Methanoperedens sp. archaea have been suggested to use a wide variety of other EAs in AOM, i.e. SO_4_^2-^, Fe^3+^ and organic compounds (Knittel and Boetius 2009; Ettwig et al. 2016; Timmers et al. 2017; Weber et al. 2017; Bai et al. 2019; Su et al. 2019). Hence, their higher abundance further supports the finding of higher AOM rates at Station 1, potentially coupled with reduction of other EAs than NO_3_^-^. In addition, ANME archaea drive SO_4_^2-^ -dependent AOM usually in syntrophic relationship with SO_4_^2-^ reducing bacterial partners (Knittel and Boetius 2009). The relative abundance of SO_4_^2-^ reducing bacteria in the depth layers representing the AOM incubation layers did not vary between the stations (Fig. S2A). However, the vertical depth patterns of *Ca*. Methanoperedens and total SO_4_^2-^ reducers (ρ = 0.81, p < 0.01) and the dominant SO_4_^2-^ reducing taxa, i.e. *Desulfarculaceae* (ρ = 0.67, p < 0.05), *Desulfobacterales* (ρ = 0.64, p < 0.05) and *Desulfobacca* (ρ = 0.71, p < 0.05,) were coherent at Station 1 (Fig. S2B), where also the highest AOM rates were observed. In contrast, such coherent patterns were not observed for Station 2 and 3. Nevertheless, these results further hint that SO_4_^2-^ - mediated AOM takes place in sediments of the study lake.

Although *Methylococcales* are not considered to be anaerobic methanotrophs, they may include species that can couple CH_4_ oxidation with reduction of NO_3_^-^/NO_2_^-^, Fe^3+^ minerals and organic EAs in hypoxic and anoxic conditions (Kits et al. 2015; Oswald et al. 2016, 2017; Martinez-Cruz et al. 2017; Rissanen et al. 2018; Zheng et al. 2020). As above for methanogens, we could not construct MAGs of *Methylococcales* or any other methanotrophic taxa, probably due to their too low abundance (see Supplementary Results and Table S1). However, according to previous MAG studies, the *Methylococcales* genera that were dominant in the study lake, i.e. *Methylobacter* and *Crenothrix* (60 – 89 % of the 16S rRNA gene amplicons affiliated to *Methylococcales* in the study lake) (Fig. S3), include members that harbored genetic potential for reduction of Fe^3+^ minerals and organic EAs in the hypoxic and anoxic water column layers of humic lakes, including closely located boreal Lake Lovojärvi (within 4 km from the study lake) (He et al. 2019; Rissanen et al. 2020). Hence, in addition to variation in abundance of *Ca*. Methanoperedens archaea, the higher abundance of *Methylococcales* also supports the observed higher AOM rates at Station 1, potentially coupled with reduction of Fe^3+^ minerals and organic EAs.

The putative NO_2_^-^ reducing AOM-driving bacteria in the genus *Ca*. Methylomirabilis had a higher relative abundance than *Ca*. Methanoperedens archaea at each station (Figs. 3 and 5). Furthermore, their relative abundance was much higher upper in the sediment profile than within the incubation layers especially at Station 2 and 3 (Fig. 3). These results suggest that NO_2_^-^ is an important EA for AOM in the study lake, as shown previously for AOM of various freshwater ecosystems, including lakes (Deutzmann et al. 2014; Welte et al. 2016).

Recent results also suggest that besides ANME archaea, methanogens may drive AOM coupled with electron transfer to solid EAs, e.g. Fe^3+^ minerals (Bar-Or et al. 2017; Yu et al. 2021). However, none of the methanogenic taxa in the AOM incubation layers had a significantly higher relative abundance at Station 1 than at other stations (Fig. S4). Thus, variations in the methanogenic community did not explain the differences in the AOM rates between stations, which suggests that methanogens did not contribute to AOM in the study lake.

## Conclusions

According to geochemical profiles as well as experimental incubations, AOM took place in the sediments of the study lake. However, despite geochemical profiles showing simultaneous consumption of CH_4_ and NO_3_^-^ in the depth layers chosen for the experimental incubations, the AOM rates were independent of NO_3_^-^ concentration. These results suggest that AOM may not be directly coupled to nitrate reduction in sediments of nitrate-rich, oligo-mesotrophic, boreal lakes. Instead, the geochemical profiles and the microbiological data both hint towards contribution of Fe^3+^ and SO_4_^2-^, while microbiological data further hint towards contribution of organic EAs and NO_2_^-^ in driving AOM in the sediments of such ecosystems, which needs to be confirmed in future experimental studies.

## Supporting information

Supplementary Results

## Code availability

Not applicable

## Author’s contributions

AJR, HJ and TJ designed the sampling and the experiments. All authors took part in acquisition, analysis and interpretation of the data. AJR drafted the manuscript, while all other authors took part in making substantial revision of the draft. All authors approved the version to be published.

## Acknowledgements

We thank staff at the Lammi Biological Station for their support in field and laboratory work.

