## Supplementary Results for "Anaerobic oxidation of methane in sediments of a nitrate-rich, oligo-mesotrophic boreal lake"

By using metagenomic binning of the assembled shotgun metagenomic sequence data, we aimed at specifically showing that methanotrophs in the sediments of the study lake have genetic potential to reduce $NO_3^-$, $Fe^{3+}$ and organic electron acceptors. Altogether, 337 bins were constructed, of which 78 were of at least of medium quality (completeness > 50 %, contamination < 10 %) (Table S1). None of the bins was affiliated with aerobic MOBs, ANME archaea or methanogenic archaea, while one low quality bin (completeness 41%) was affiliated with *Ca*. Methylomirabilis sp (Table S1). Furthermore, genes coding for methyl coenzyme M reductase, the key enzyme in methanogenesis and in ANME archaea - mediated AOM, were not found in either of the bins. In addition, genes coding for particulate (pmoCAB) or soluble methane monooxygenase (mmoXYBZDC) of aerobic MOBs were found only incompletely (i.e. either pmoA, pmoB or mmoC) and only from 2 medium quality bins and 7 low quality bins. Thus, despite their activity, both aerobic and anaerobic methanotrophs very likely had too low abundance to be detected by the metagenomic binning analysis, and hence, this data is not considered further in this study. However, this data will be presented in more detail in our subsequent paper on Fe and N cycling reactions in the study lake (Jäntti et al. in prep.).
