## Supplementary Figures for "Anaerobic oxidation of methane in sediments of a nitrate-rich, oligo-mesotrophic boreal lake"

### Supplementary Figures S1-S4

#### Anaerobic oxidation of methane in sediments of a nitrate-rich, oligo-mesotrophic, boreal lake

Antti J Rissanen[1,*], Tom Jilbert[2], Asko Simojoki[3], Rahul Mangayil[1], Sanni L Aalto[4,5,a], Sari Peura[6], Helena Jäntti[4]

[1]Faculty of Engineering and Natural Sciences, Tampere University, Korkeakoulunkatu 6, FI-33720, Tampere, Finland

[2]Ecosystems and Environment Research Program, Faculty of Biological and Environmental Sciences, University of Helsinki, P.O. Box 65, FI-00014, Helsinki, Finland

[3]Department of Agricultural Sciences (Environmental Soil Science), Faculty of Agriculture and Forestry, University of Helsinki, P.O. Box 56, FI-00014, Helsinki, Finland

[4]Department of Environmental and Biological Sciences, University of Eastern Finland, Yliopistonranta 1 E, FI-70210, Kuopio, Finland

[5]Department of Biological and Environmental Sciences, University of Jyväskylä, Survontie 9 C, FI-40014, Jyväskylä, Finland

[6]Department of Forest Mycology and Plant Pathology, Science for Life Laboratory, Swedish University of Agricultural Sciences, Almas allé 5, SE-75651, Uppsala, Sweden

[a]Current address: Technical University of Denmark, DTU Aqua, Section for Aquaculture, The North Sea Research Centre, P.O. Box 101, DK-9850, Hirtshals, Denmark

*Corresponding author: Antti J Rissanen, Faculty of Engineering and Natural Sciences, Tampere University, Korkeakoulunkatu 6, FI-33720, Tampere, Finland. Tel: +358 40 1981145; Fax: +358 3 3641392;

#### Fig. S1

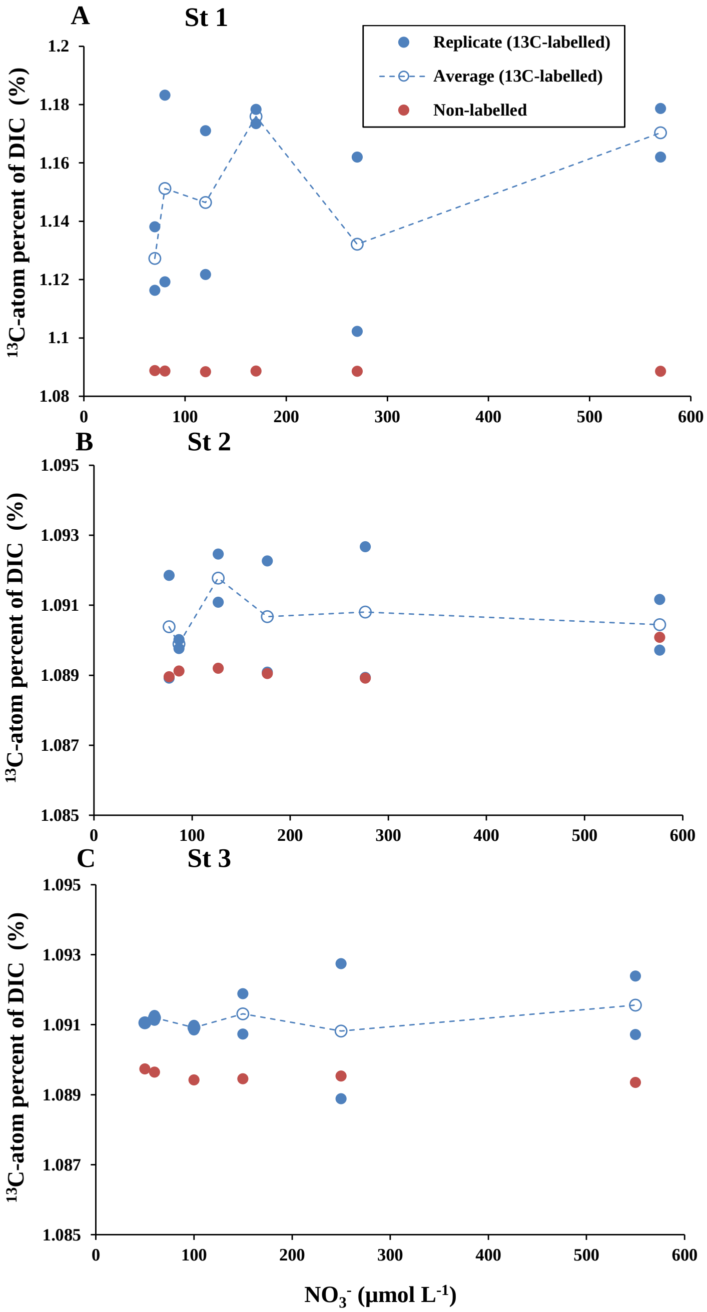


**Fig. S1**. $^{13}C$-atom percent of DIC in $^{13}C$-labelled and non-labelled samples at different concentrations of $NO_3^-$ at **A** Station 1, **B** Station 2 and **C** Station 3. For $^{13}C$-labelled samples, values are represented as replicate and average values (n=2 at each concentration level of $NO_3^-$), while one replicate was analysed for non-labelled samples. The $NO_3^-$ concentration levels at the x-axis represent the sum of added $NO_3^-$ (i.e. 0, 10, 50, 100, 200 or 500 µmol $L^{-1}$) and $NO_3^-$ already present in slurry at the beginning of incubations (i.e. 70, 77 and 50 µmol $L^{-1}$ for stations 1, 2 and 3, respectively). No positive relationship between $NO_3^-$ and $^{13}C$-atom percent of DIC was observed at the stations. Note different scale in y-axis in **A** compared to **B** and **C**

**Fig. S2**

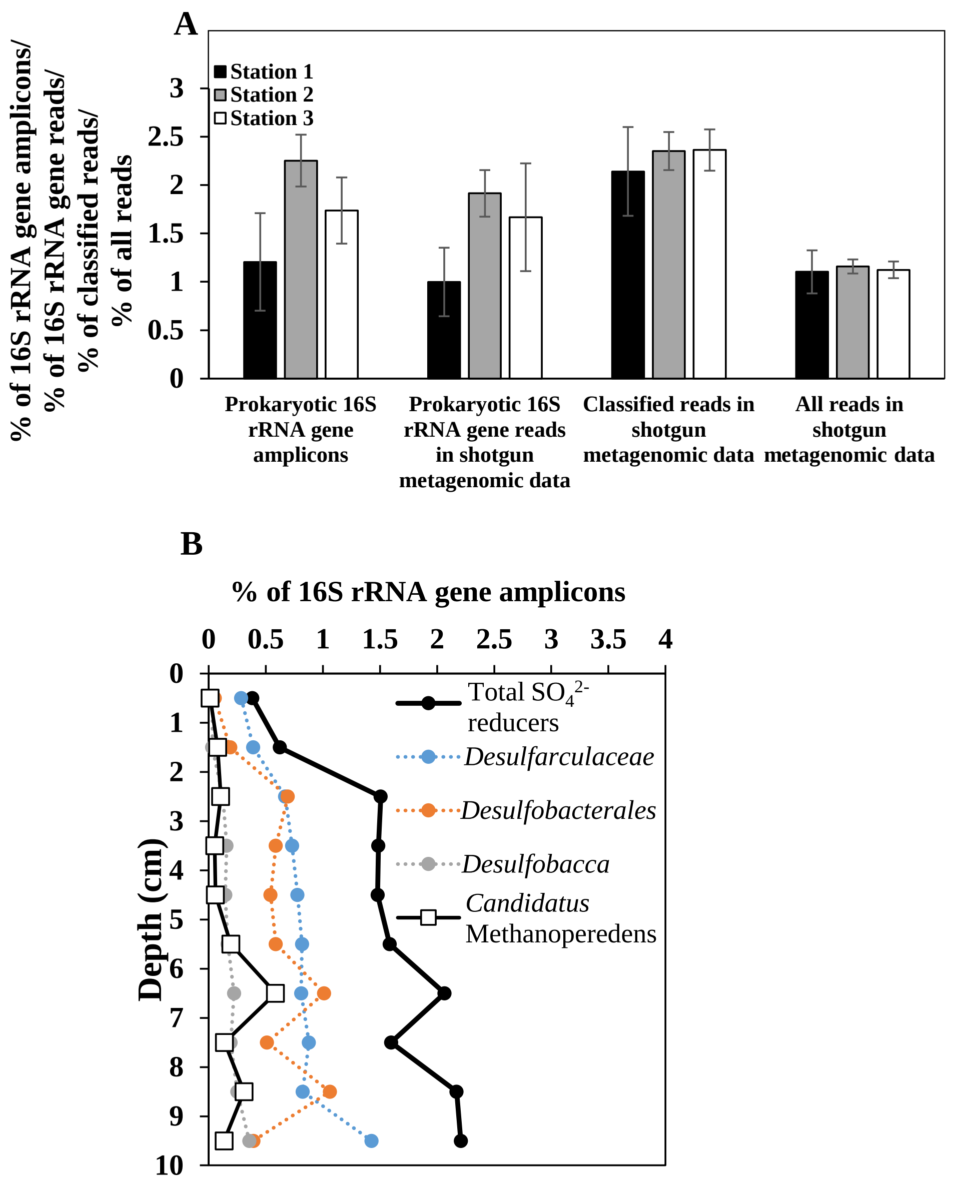


**Fig. S2**. **A** Relative abundance of total $SO_4^{2-}$ reducing bacteria in the depth layers representing the incubation layers based on 16S rRNA gene amplicon sequencing, as well as shotgun metagenomic sequencing, i.e 16S rRNA gene reads, KAIJU-classified reads expressed as relative to reads that were classified at least to domain-level, and KAIJU-classified reads expressed as relative to all reads, as well as, **B** relative abundance of total $SO_4^{2-}$ reducing bacteria, dominant $SO_4^{2-}$ reducing bacterial taxa and putative AOM-driving *Ca*. Methanoperedens sp. archaea in different depth layers at Station 1 based on 16S rRNA gene amplicon sequencing. Data is represented as an average and standard deviation of three layer-specific samples (in **A**). No significant differences were observed between different stations in RB-ANOVA analyses (in **A**)

**Fig. S3**

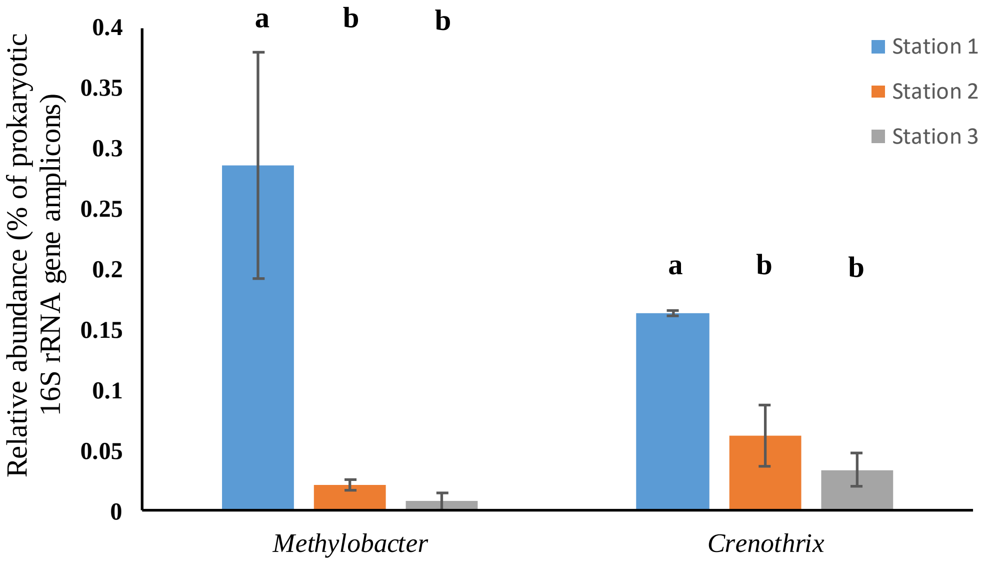


**Fig. S3**. Relative abundance of dominant genera of *Methylococcales* in the depth layers representing the incubation layers based on 16S rRNA gene amplicon sequencing. Data is represented as an average and standard deviation of three layer-specific samples. Different letters above bars indicate significant differences in RB-ANOVA and subsequent pairwise tests ($p < 0.05$)

**Fig. S4**

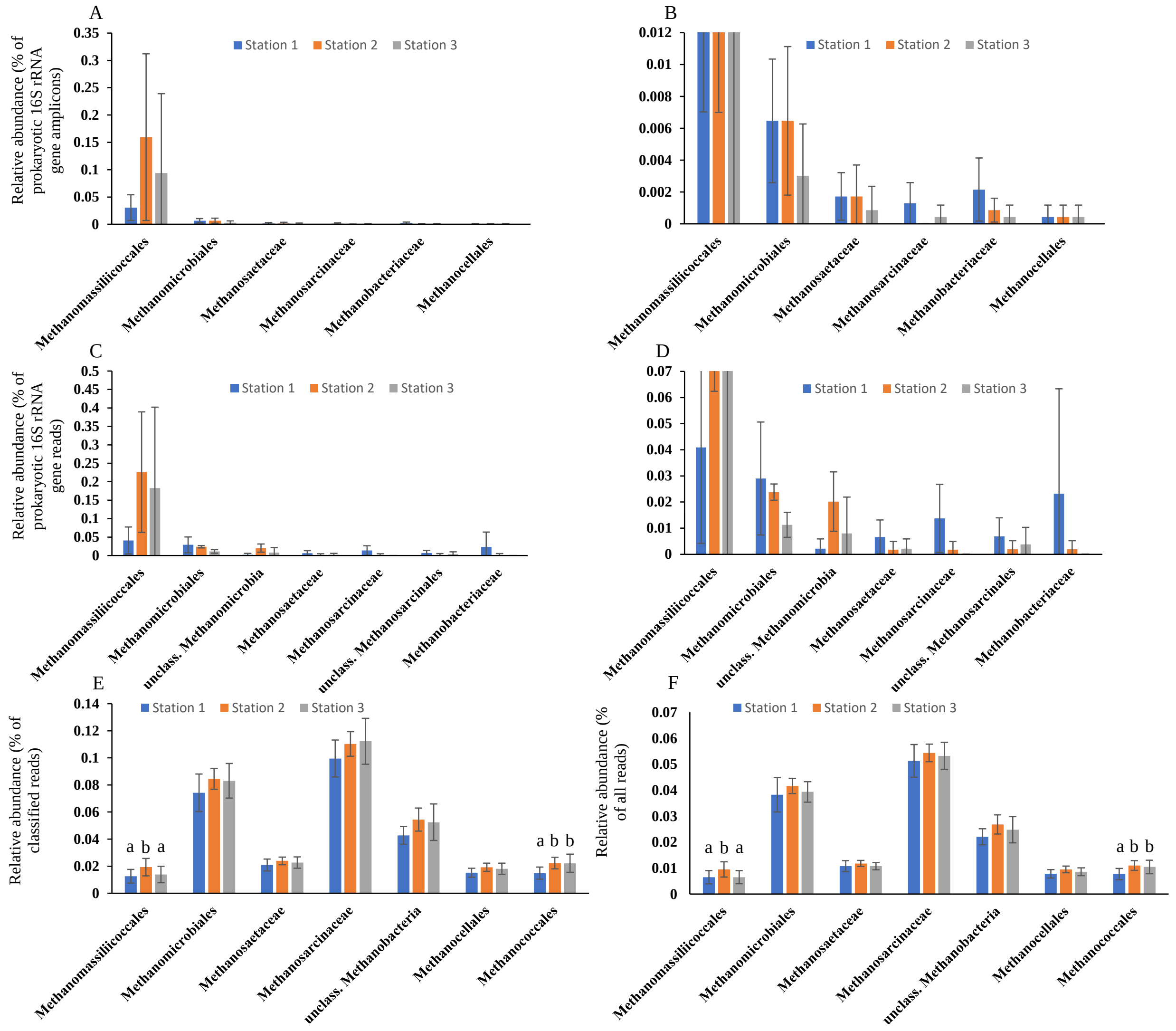


**Fig. S4**. Relative abundance of total methanogens and different methanogenic taxa in the depth layers representing the incubation layers based on **A-B** 16S rRNA gene amplicon sequencing, as well as **C-F** shotgun metagenomic sequencing, i.e. **C-D** 16S rRNA gene reads, **E** KAIJU-classified reads as relative to reads that were classified at least to domain-level and **F** KAIJU-classified reads as relative to all reads. To show the results clearly for less abundant taxa, **B** and **D** represent the same data as in **A** and **C**, respectively, but with different scaling. Data is represented as an average and standard deviation of three layer-specific samples. Different letters above bars indicate significant differences in RB-ANOVA and subsequent pairwise tests ($p < 0.05$)
