## Supplementary TableS1 for "Anaerobic oxidation of methane in sediments of a nitrate-rich, oligo-mesotrophic boreal lake"

### Supplementary Table S1

#### Anaerobic oxidation of methane in sediments of a nitrate-rich, oligo-mesotrophic, boreal lake

Antti J Rissanen[1,*], Tom Jilbert[2], Asko Simojoki[3], Rahul Mangayil[1], Sanni L Aalto[4,5,a], Sari Peura[6], Helena Jäntti[4]

[1]Faculty of Engineering and Natural Sciences, Tampere University, Korkeakoulunkatu 6, FI-33720, Tampere, Finland

[2]Ecosystems and Environment Research Program, Faculty of Biological and Environmental Sciences, University of Helsinki, P.O. Box 65, FI-00014, Helsinki, Finland

[3]Department of Agricultural Sciences (Environmental Soil Science), Faculty of Agriculture and Forestry, University of Helsinki, P.O. Box 56, FI-00014, Helsinki, Finland

[4]Department of Environmental and Biological Sciences, University of Eastern Finland, Yliopistonranta 1 E, FI-70210, Kuopio, Finland

[5]Department of Biological and Environmental Sciences, University of Jyväskylä, Survontie 9 C, FI-40014, Jyväskylä, Finland

[6]Department of Forest Mycology and Plant Pathology, Science for Life Laboratory, Swedish University of Agricultural Sciences, Almas allé 5, SE-75651, Uppsala, Sweden

[a]Current address: Technical University of Denmark, DTU Aqua, Section for Aquaculture, The North Sea Research Centre, P.O. Box 101, DK-9850, Hirtshals, Denmark

*Corresponding author: Antti J Rissanen, Faculty of Engineering and Natural Sciences, Tampere University, Korkeakoulunkatu 6, FI-33720, Tampere, Finland. Tel: +358 40 1981145; Fax: +358 3 3641392;

**TABLE S1**. Size, completeness, contamination, CH4 oxidation genes and taxonomic classification of MAGs.
Medium and good quality MAGs (i.e. completeness > 50%, contamination < 10%) are denoted as qual. class 1, other MAGs are qual. class 2.

| Bin no. | Size (mb) | Comp. (%) | Cont. (%) | Qual. class | CH4 ox. gene | Taxonomic classification based on CAT/BAT - tool. |
|---|---|---|---|---|---|---|
| 249 | 2.72 | 76.72 | 0 | 1 | | Bacteria : 1.00unclassified Bacteria (no rank): 0.78Bacteria candidate phyla (no rank): 0.77Candidatus Eisenbacteria (phylum): 0.76 |
| 309 | 0.77 | 72.43 | 0 | 1 | | Archaea : 0.74DPANN group (no rank): 0.32 |
| 334 | 1.01 | 65.07 | 0.07 | 1 | mmoC | Bacteria : 0.99unclassified Bacteria (no rank): 0.56Bacteria candidate phyla (no rank): 0.54Patescibacteria group (no rank): 0.54Microgenomates group (no rank): 0.52Candidatus Roizmanbacteria (phylum): 0.43 |
| 181 | 1.83 | 89.55 | 0.2 | 1 | | Bacteria : 0.95Terrabacteria group (no rank): 0.75Chloroflexi (phylum): 0.73 |
| 196 | 1.02 | 66.85 | 0.32 | 1 | | Archaea : 0.99TACK group (no rank): 0.92Thaumarchaeota (phylum): 0.91unclassified Thaumarchaeota (no rank): 0.82unclassified Thaumarchaeota (miscellaneous) (no rank): 0.81Thaumarchaeota archaeon CSP1-1 (species): 0.80 |
| 289 | 1.49 | 50.9 | 0.48 | 1 | | Bacteria : 0.98FCB group (no rank): 0.83Bacteroidetes/Chlorobi group (no rank): 0.83Bacteroidetes (phylum): 0.81unclassified Bacteroidetes (no rank): 0.46unclassified Bacteroidetes (miscellaneous) (no rank): 0.46 |
| 203 | 1.88 | 97.44 | 0.69 | 1 | | Bacteria : 0.99Proteobacteria (phylum): 0.93Betaproteobacteria (class): 0.74 |
| 117 | 0.84 | 67.43 | 0.99 | 1 | | Bacteria : 0.97Terrabacteria group (no rank): 0.83Chloroflexi (phylum): 0.83Dehalococcoidia (class): 0.55unclassified Dehalococcoidia (no rank): 0.54Dehalococcoidia bacterium (species): 0.54 |
| 191 | 1.8 | 65.19 | 1.07 | 1 | | Bacteria : 0.99unclassified Bacteria (no rank): 0.45Bacteria candidate phyla (no rank): 0.45candidate division NC10 (phylum): 0.37unclassified candidate division NC10 (no rank): 0.36 |
| 296 | 0.81 | 50.55 | 1.08 | 1 | | Bacteria : 0.99PVC group (no rank): 0.93Candidatus Omnitrophica (phylum): 0.93unclassified Candidatus Omnitrophica (no rank): 0.93 |
| 169 | 2.94 | 68.01 | 1.09 | 1 | | Bacteria : 0.99Proteobacteria (phylum): 0.77delta/epsilon subdivisions (subphylum): 0.73Deltaproteobacteria (class): 0.73unclassified Deltaproteobacteria (no rank): 0.72unclassified Deltaproteobacteria (miscellaneous) (no rank): 0.72Deltapr |
| 97 | 3 | 73.62 | 1.14 | 1 | | Bacteria : 0.92PVC group (no rank): 0.30 |
| 286 | 1.38 | 55.87 | 1.51 | 1 | | Bacteria : 0.93Proteobacteria (phylum): 0.34delta/epsilon subdivisions (subphylum): 0.32Deltaproteobacteria (class): 0.32 |
| 231 | 0.89 | 64.69 | 1.69 | 1 | mmoC | Bacteria : 0.92unclassified Bacteria (no rank): 0.78Bacteria candidate phyla (no rank): 0.76Candidatus Dojkabacteria (no rank): 0.59 |
| 153 | 0.6 | 56.31 | 1.87 | 1 | | Archaea : 0.80DPANN group (no rank): 0.40Candidatus Woesearchaeota (phylum): 0.32 |
| 37 | 0.84 | 60.36 | 1.98 | 1 | | Bacteria : 0.80 |
| 39 | 0.76 | 59.9 | 1.98 | 1 | | Bacteria : 0.99unclassified Bacteria (no rank): 0.74Bacteria candidate phyla (no rank): 0.74Patescibacteria group (no rank): 0.72Microgenomates group (no rank): 0.69 |
| 319 | 4.82 | 92.9 | 1.99 | 1 | | Bacteria : 0.99Proteobacteria (phylum): 0.69delta/epsilon subdivisions (subphylum): 0.65Deltaproteobacteria (class): 0.65unclassified Deltaproteobacteria (no rank): 0.37unclassified Deltaproteobacteria (miscellaneous) (no rank): 0.37Deltapr |
| 127 | 1.27 | 78.41 | 2.1 | 1 | | Bacteria : 1.00Proteobacteria (phylum): 0.97delta/epsilon subdivisions (subphylum): 0.96Deltaproteobacteria (class): 0.96unclassified Deltaproteobacteria (no rank): 0.93unclassified Deltaproteobacteria (miscellaneous) (no rank): 0.93 |
| 323 | 2.36 | 62.38 | 2.16 | 1 | | Bacteria : 0.99Proteobacteria (phylum): 0.84Alphaproteobacteria (class): 0.73Rhodospirillales (order): 0.53Rhodospirillaceae (family): 0.47unclassified Rhodospirillaceae (no rank): 0.37 |
| 298 | 3.73 | 88.01 | 2.23 | 1 | | Bacteria : 1.00FCB group (no rank): 0.76Bacteroidetes/Chlorobi group (no rank): 0.76Bacteroidetes (phylum): 0.75 |
| 253 | 2.7 | 67.14 | 2.25 | 1 | | Bacteria : 0.97PVC group (no rank): 0.32 |
| 274 | 2.72 | 77.15 | 2.31 | 1 | | Bacteria : 0.99Proteobacteria (phylum): 0.88Gammaproteobacteria (class): 0.69Chromatiales (order): 0.59unclassified Chromatiales (no rank): 0.59unclassified Chromatiales (miscellaneous) (no rank): 0.59Chromatiales bacterium (species): 0.59 |
| 67 | 2.11 | 90.67 | 2.44 | 1 | | Bacteria : 0.99Proteobacteria (phylum): 0.90Gammaproteobacteria (class): 0.80unclassified Gammaproteobacteria (no rank): 0.72unclassified Gammaproteobacteria (miscellaneous) (no rank): 0.71 |
| 167 | 2.6 | 65.32 | 2.58 | 1 | | Bacteria : 0.99Proteobacteria (phylum): 0.89delta/epsilon subdivisions (subphylum): 0.89Deltaproteobacteria (class): 0.89unclassified Deltaproteobacteria (no rank): 0.84unclassified Deltaproteobacteria (miscellaneous) (no rank): 0.84 |
| 244 | 2.93 | 89.5 | 2.6 | 1 | | Bacteria : 0.98unclassified Bacteria (no rank): 0.75unclassified Bacteria (miscellaneous) (no rank): 0.74bacterium (species): 0.74 |
| 134 | 2.22 | 73.67 | 2.73 | 1 | | Bacteria : 0.98Nitrospirae (phylum): 0.37 |
| 156 | 1.3 | 85.6 | 2.79 | 1 | | Bacteria : 1.00PVC group (no rank): 0.66Candidatus Omnitrophica (phylum): 0.66unclassified Candidatus Omnitrophica (no rank): 0.66Omnitrophica WOR_2 bacterium SM23_72 (species): 0.37 |
| 14 | 1.38 | 69.48 | 2.8 | 1 | | Archaea : 0.91TACK group (no rank): 0.86Candidatus Bathyarchaeota (phylum): 0.85unclassified Candidatus Bathyarchaeota (no rank): 0.73Candidatus Bathyarchaeota archaeon (species): 0.67 |
| 277 | 3.72 | 61.59 | 2.9 | 1 | | Bacteria : 0.99Proteobacteria (phylum): 0.70Gammaproteobacteria (class): 0.49unclassified Gammaproteobacteria (no rank): 0.40unclassified Gammaproteobacteria (miscellaneous) (no rank): 0.35Gammaproteobacteria bacterium (species): |
| 85 | 1.53 | 100 | 2.91 | 1 | | Archaea : 1.00TACK group (no rank): 0.94Thaumarchaeota (phylum): 0.94unclassified Thaumarchaeota (no rank): 0.42Candidatus Nitrosotalea (genus): 0.30 |
| 78 | 1.31 | 67.53 | 3.13 | 1 | | Bacteria : 0.98PVC group (no rank): 0.34Candidatus Omnitrophica (phylum): 0.33unclassified Candidatus Omnitrophica (no rank): 0.33 |
| 124 | 2.21 | 71.05 | 3.23 | 1 | | Bacteria : 0.99FCB group (no rank): 0.94Bacteroidetes/Chlorobi group (no rank): 0.94Bacteroidetes (phylum): 0.93unclassified Bacteroidetes (no rank): 0.80unclassified Bacteroidetes (miscellaneous) (no rank): 0.80 |
| 160 | 2.29 | 50.75 | 3.26 | 1 | | Bacteria : 0.97Proteobacteria (phylum): 0.72delta/epsilon subdivisions (subphylum): 0.70Deltaproteobacteria (class): 0.70 |
| 75 | 0.44 | 63.19 | 3.27 | 1 | | Archaea : 0.82 |
| 4 | 2 | 56.41 | 3.48 | 1 | | Bacteria : 0.99FCB group (no rank): 0.63Gemmatimonadetes (phylum): 0.62 |
| 140 | 3.17 | 91.48 | 3.57 | 1 | | Bacteria : 1.00Proteobacteria (phylum): 0.86Betaproteobacteria (class): 0.66unclassified Betaproteobacteria (no rank): 0.55unclassified Betaproteobacteria (miscellaneous) (no rank): 0.55 |
| 143 | 2.72 | 56.99 | 3.61 | 1 | | Bacteria : 0.99unclassified Bacteria (no rank): 0.43unclassified Bacteria (miscellaneous) (no rank): 0.41bacterium (species): 0.41 |
| 202 | 1.54 | 50.6 | 3.67 | 1 | | Bacteria : 0.96Terrabacteria group (no rank): 0.72Chloroflexi (phylum): 0.69unclassified Chloroflexi (no rank): 0.46unclassified Chloroflexi (miscellaneous) (no rank): 0.45 |
| 209 | 0.78 | 69.17 | 3.76 | 1 | | Bacteria : 0.94unclassified Bacteria (no rank): 0.73Bacteria candidate phyla (no rank): 0.66Patescibacteria group (no rank): 0.63Parcubacteria group (no rank): 0.59 |
| 282 | 1.08 | 63.22 | 3.78 | 1 | | Archaea : 0.97Euryarchaeota (phylum): 0.89unclassified Euryarchaeota (no rank): 0.68unclassified Euryarchaeota (miscellaneous) (no rank): 0.68 |
| 46 | 2.62 | 84.13 | 3.83 | 1 | | Bacteria : 0.98FCB group (no rank): 0.64Bacteroidetes/Chlorobi group (no rank): 0.60Ignavibacteriae (phylum): 0.33 |
| 241 | 3.16 | 88.07 | 3.98 | 1 | | Bacteria : 0.97PVC group (no rank): 0.64Planctomycetes (phylum): 0.61unclassified Planctomycetes (no rank): 0.33 |
| 114 | 5.96 | 84.4 | 4.05 | 1 | | Bacteria : 0.99Acidobacteria (phylum): 0.63unclassified Acidobacteria (no rank): 0.57unclassified Acidobacteria (miscellaneous) (no rank): 0.57Acidobacteria bacterium (species): 0.54 |
| 96 | 3.38 | 72.73 | 4.09 | 1 | | Bacteria : 0.98PVC group (no rank): 0.69Planctomycetes (phylum): 0.66Phycisphaerae (class): 0.38unclassified Phycisphaerae (no rank): 0.35Phycisphaerae bacterium RAS1 (species): 0.34 |
| 276 | 2.32 | 50.56 | 4.27 | 1 | | Bacteria : 0.98Acidobacteria (phylum): 0.56unclassified Acidobacteria (no rank): 0.52unclassified Acidobacteria (miscellaneous) (no rank): 0.52Acidobacteria bacterium (species): 0.51 |
| 86 | 3.37 | 97.8 | 4.4 | 1 | | Bacteria : 0.99FCB group (no rank): 0.65Gemmatimonadetes (phylum): 0.64Gemmatimonadetes (class): 0.30Gemmatimonadales* (order): 0.30 |
| 26 | 2.68 | 86.58 | 4.5 | 1 | | Bacteria : 1.00FCB group (no rank): 0.71Gemmatimonadetes (phylum): 0.71unclassified Gemmatimonadetes (no rank): 0.45Gemmatimonadetes bacterium (species): 0.39 |
| 210 | 2.63 | 64.97 | 4.93 | 1 | | Archaea : 0.95TACK group (no rank): 0.92Thaumarchaeota (phylum): 0.91Nitrosopumilales (order): 0.67Nitrosopumilaceae (family): 0.35 |
| 91 | 1.69 | 73.77 | 5.07 | 1 | | Bacteria : 0.98Proteobacteria (phylum): 0.75 |
| 119 | 2.57 | 57.05 | 5.17 | 1 | | Bacteria : 0.99FCB group (no rank): 0.69Gemmatimonadetes (phylum): 0.69unclassified Gemmatimonadetes (no rank): 0.63 |
| 184 | 1.1 | 77.57 | 5.22 | 1 | | Archaea : 0.90Euryarchaeota (phylum): 0.73unclassified Euryarchaeota (no rank): 0.71unclassified Euryarchaeota (miscellaneous) (no rank): 0.71Euryarchaeota archaeon SM23-78 (species): 0.71 |
| 182 | 1.77 | 78.05 | 5.25 | 1 | | Bacteria : 0.99Proteobacteria (phylum): 0.78 |
| 145 | 3.21 | 93.45 | 5.43 | 1 | | Bacteria : 1.00Proteobacteria (phylum): 0.95Alphaproteobacteria (class): 0.50 |
| 116 | 3.64 | 84.92 | 5.49 | 1 | | Bacteria : 0.99FCB group (no rank): 0.61Gemmatimonadetes (phylum): 0.60 |
| 90 | 1.8 | 85.26 | 5.56 | 1 | | Bacteria : 0.97Terrabacteria group (no rank): 0.85Cyanobacteria/Melainabacteria group (no rank): 0.84Candidatus Saganbacteria (no rank): 0.84 |

| | | | | | |
|---|---|---|---|---|---|
| 175 | 2.52 | 96.59 | 5.91 | 1 | Bacteria : 0.98Nitrospirae (phylum): 0.81unclassified Nitrospirae (no rank): 0.40 |
| 3 | 3.25 | 64.77 | 6.27 | 1 | Bacteria : 0.99FCB group (no rank): 0.92Bacteroidetes/Chlorobi group (no rank): 0.92Bacteroidetes (phylum): 0.91unclassified Bacteroidetes (no rank): 0.30unclassified Bacteroidetes (miscellaneous) (no rank): 0.30 |
| 98 | 3.22 | 75.99 | 6.54 | 1 | Bacteria : 0.99PVC group (no rank): 0.82Verrucomicrobia (phylum): 0.80Opitutae (class): 0.45 |
| 50 | 1.79 | 74.28 | 6.82 | 1 | Bacteria : 1.00Nitrospirae (phylum): 0.92unclassified Nitrospirae (no rank): 0.77 |
| 103 | 8.6 | 90.52 | 6.85 | 1 | Bacteria : 0.99PVC group (no rank): 0.55Planctomycetes (phylum): 0.49 |
| 295 | 4.19 | 92.02 | 6.86 | 1 | Bacteria : 0.99Proteobacteria (phylum): 0.87Betaproteobacteria (class): 0.77unclassified Betaproteobacteria (no rank): 0.73unclassified Betaproteobacteria (miscellaneous) (no rank): 0.73Betaproteobacteria bacterium (species): 0.54 |
| 149 | 3.45 | 95.42 | 6.96 | 1 | Bacteria : 0.99FCB group (no rank): 0.66Gemmatimonadetes (phylum): 0.65unclassified Gemmatimonadetes (no rank): 0.60 |
| 245 | 1.42 | 57.57 | 7.18 | 1 | Bacteria : 0.99Terrabacteria group (no rank): 0.84Actinobacteria (phylum): 0.81Actinobacteria (class): 0.37unclassified Actinobacteria (class) (no rank): 0.35unclassified Actinobacteria (class) (miscellaneous) (no rank): 0.35Actinobacteria bacterium (species): 0.33 |
| 147 | 4.02 | 85.33 | 7.42 | 1 | Bacteria : 0.99Proteobacteria (phylum): 0.75delta/epsilon subdivisions (subphylum): 0.72Deltaproteobacteria (class): 0.72unclassified Deltaproteobacteria (no rank): 0.69unclassified Deltaproteobacteria (miscellaneous) (no rank): 0.69Deltapr |
| 188 | 2.84 | 73.11 | 7.76 | 1 | Bacteria : 0.97FCB group (no rank): 0.70Bacteroidetes/Chlorobi group (no rank): 0.68Ignavibacteriae (phylum): 0.55unclassified Ignavibacteriae (no rank): 0.45Ignavibacteriae bacterium (species): 0.45 |
| 310 | 2.06 | 57.16 | 7.98 | 1 | Bacteria : 1.00Proteobacteria (phylum): 0.86delta/epsilon subdivisions (subphylum): 0.84Deltaproteobacteria (class): 0.84unclassified Deltaproteobacteria (no rank): 0.84unclassified Deltaproteobacteria (miscellaneous) (no rank): 0.84Deltapr |
| 256 | 1 | 51.58 | 8.34 | 1 | Bacteria : 0.98PVC group (no rank): 0.38Candidatus Omnitrophica (phylum): 0.38unclassified Candidatus Omnitrophica (no rank): 0.38 |
| 41 | 3.29 | 93.39 | 8.39 | 1 | Bacteria : 0.99Proteobacteria (phylum): 0.78 |
| 33 | 4.09 | 90.66 | 8.41 | 1 | Bacteria : 0.98FCB group (no rank): 0.87Bacteroidetes/Chlorobi group (no rank): 0.87Bacteroidetes (phylum): 0.85unclassified Bacteroidetes (no rank): 0.63unclassified Bacteroidetes (miscellaneous) (no rank): 0.63Bacteroidetes bacterium (sp |
| 315 | 3.2 | 80.49 | 8.76 | 1 | Bacteria : 0.99FCB group (no rank): 0.62Bacteroidetes/Chlorobi group (no rank): 0.60Ignavibacteriae (phylum): 0.36Ignavibacteria (class): 0.31 |
| 121 | 0.81 | 50.01 | 8.81 | 1 | Bacteria : 0.98unclassified Bacteria (no rank): 0.82Bacteria candidate phyla (no rank): 0.80Patescibacteria group (no rank): 0.31 |
| 20 | 1.83 | 84.26 | 9.02 | 1 | Bacteria : 0.74 |
| 281 | 3.87 | 92.48 | 9.04 | 1 | Bacteria : 0.99unclassified Bacteria (no rank): 0.77Bacteria candidate phyla (no rank): 0.76Candidatus Eisenbacteria (phylum): 0.72 |
| 270 | 2.37 | 67.18 | 9.04 | 1 | Bacteria : 0.99Terrabacteria group (no rank): 0.89Chloroflexi (phylum): 0.89 |
| 95 | 5.8 | 69.74 | 9.21 | 1 | Bacteria : 0.99Proteobacteria (phylum): 0.75delta/epsilon subdivisions (subphylum): 0.71Deltaproteobacteria (class): 0.71unclassified Deltaproteobacteria (no rank): 0.70unclassified Deltaproteobacteria (miscellaneous) (no rank): 0.70Deltapr |
| 314 | 3.21 | 69.35 | 9.76 | 1 | Bacteria : 0.98Acidobacteria (phylum): 0.53unclassified Acidobacteria (no rank): 0.49 |
| 330 | 3.33 | 94.51 | 9.99 | 1 | Bacteria : 1.00unclassified Bacteria (no rank): 0.82Bacteria candidate phyla (no rank): 0.81Candidatus Eisenbacteria (phylum): 0.79Candidatus Eisenbacteria bacterium (species): 0.57 |
| 322 | 5.19 | 95.73 | 10 | 2 | Bacteria : 0.99Acidobacteria (phylum): 0.85unclassified Acidobacteria (no rank): 0.79unclassified Acidobacteria (miscellaneous) (no rank): 0.79Acidobacteria bacterium (species): 0.64 |
| 25 | 0.66 | 50.05 | 10.29 | 2 | Bacteria : 0.98unclassified Bacteria (no rank): 0.89Bacteria candidate phyla (no rank): 0.88Patescibacteria group (no rank): 0.87Microgenomates group (no rank): 0.83 |
| 92 | 2.85 | 98.18 | 10.56 | 2 | Bacteria : 0.98Nitrospirae (phylum): 0.69 |
| 154 | 2.67 | 59.91 | 10.95 | 2 | Bacteria : 0.98Proteobacteria (phylum): 0.65Alphaproteobacteria (class): 0.48 |
| 18 | 6.68 | 81.14 | 11.05 | 2 | Bacteria : 0.96unclassified Bacteria (no rank): 0.57 |
| 5 | 2.11 | 51.34 | 11.38 | 2 | Bacteria : 0.99Proteobacteria (phylum): 0.74delta/epsilon subdivisions (subphylum): 0.72Deltaproteobacteria (class): 0.72Syntrophobacterales (order): 0.36 |
| 111 | 2.13 | 53.15 | 11.54 | 2 | Bacteria : 0.98Acidobacteria (phylum): 0.65unclassified Acidobacteria (no rank): 0.49unclassified Acidobacteria (miscellaneous) (no rank): 0.49Acidobacteria bacterium (species): 0.44 |
| 157 | 0.72 | 53.19 | 11.74 | 2 | Bacteria : 0.99unclassified Bacteria (no rank): 0.75Bacteria candidate phyla (no rank): 0.72Patescibacteria group (no rank): 0.71Parcubacteria group (no rank): 0.69Candidatus Moranbacteria (phylum): 0.64 |
| 317 | 3.34 | 93.99 | 12.19 | 2 | Bacteria : 0.99FCB group (no rank): 0.72Bacteroidetes/Chlorobi group (no rank): 0.70Ignavibacteriae (phylum): 0.54Ignavibacteria (class): 0.50unclassified Ignavibacteria (no rank): 0.49 |
| 8 | 6.19 | 82.01 | 12.64 | 2 | Bacteria : 0.99Proteobacteria (phylum): 0.75Betaproteobacteria (class): 0.37 |
| 47 | 3.8 | 75.34 | 12.93 | 2 | Bacteria : 1.00unclassified Bacteria (no rank): 0.69Bacteria candidate phyla (no rank): 0.68Candidatus Eisenbacteria (phylum): 0.65 |
| 297 | 4.75 | 86.96 | 13.46 | 2 | Bacteria : 0.99Proteobacteria (phylum): 0.72Alphaproteobacteria (class): 0.50 |
| 185 | 1.65 | 72.72 | 15.4 | 2 | Bacteria : 0.98PVC group (no rank): 0.83Candidatus Omnitrophica (phylum): 0.83unclassified Candidatus Omnitrophica (no rank): 0.83 |
| 32 | 4.88 | 66.1 | 15.48 | 2 | Bacteria : 0.99Acidobacteria (phylum): 0.74unclassified Acidobacteria (no rank): 0.54unclassified Acidobacteria (miscellaneous) (no rank): 0.54Acidobacteria bacterium (species): 0.36 |
| 165 | 5.19 | 58.62 | 15.52 | 2 | Bacteria : 0.99unclassified Bacteria (no rank): 0.41unclassified Bacteria (miscellaneous) (no rank): 0.38bacterium (species): 0.38 |
| 335 | 3.81 | 80.18 | 15.64 | 2 | Bacteria : 0.99PVC group (no rank): 0.65Planctomycetes (phylum): 0.64unclassified Planctomycetes (no rank): 0.62Planctomycetes bacterium (species): 0.38 |
| 194 | 2.68 | 50.92 | 15.93 | 2 | Bacteria : 1.00FCB group (no rank): 0.77Gemmatimonadetes (phylum): 0.77unclassified Gemmatimonadetes (no rank): 0.71 |
| 237 | 1.09 | 76.56 | 15.95 | 2 | Bacteria : 0.96unclassified Bacteria (no rank): 0.71Bacteria candidate phyla (no rank): 0.70Patescibacteria group (no rank): 0.67Microgenomates group (no rank): 0.57Candidatus Levybacteria (phylum): 0.37 |
| 333 | 2.48 | 63.01 | 19.25 | 2 | Bacteria : 0.99FCB group (no rank): 0.69Gemmatimonadetes (phylum): 0.69Gemmatimonadetes (class): 0.49Gemmatimonadales (order): 0.49unclassified Gemmatimonadales (no rank): 0.48Gemmatimonadales bacterium* (species): 0.48 |
| 265 | 2.63 | 81.55 | 20.11 | 2 | Bacteria : 0.92Proteobacteria (phylum): 0.36delta/epsilon subdivisions (subphylum): 0.35Deltaproteobacteria (class): 0.34 |
| 101 | 1.65 | 57.9 | 20.87 | 2 | Archaea : 0.98TACK group (no rank): 0.97Thaumarchaeota (phylum): 0.96Nitrosopumilales (order): 0.75 |
| 40 | 5.17 | 68.1 | 23.3 | 2 | Bacteria : 1.00Proteobacteria (phylum): 0.85delta/epsilon subdivisions (subphylum): 0.81Deltaproteobacteria (class): 0.81unclassified Deltaproteobacteria (no rank): 0.81unclassified Deltaproteobacteria (miscellaneous) (no rank): 0.81Deltapr |
| 328 | 4.44 | 82.78 | 23.83 | 2 | Bacteria : 1.00FCB group (no rank): 0.68Gemmatimonadetes (phylum): 0.67unclassified Gemmatimonadetes (no rank): 0.44Gemmatimonadetes bacterium (species): 0.35 |
| 178 | 4.46 | 67.08 | 24.53 | 2 | Bacteria : 0.99Terrabacteria group (no rank): 0.93Chloroflexi (phylum): 0.91 |
| 195 | 5.07 | 74.63 | 25.36 | 2 | Bacteria : 0.99Proteobacteria (phylum): 0.60 |
| 201 | 8.71 | 70.87 | 25.92 | 2 | Bacteria : 0.97 |
| 21 | 3.47 | 76.24 | 27.52 | 2 | Bacteria : 0.99FCB group (no rank): 0.88Bacteroidetes/Chlorobi group (no rank): 0.88Bacteroidetes (phylum): 0.85unclassified Bacteroidetes (no rank): 0.61unclassified Bacteroidetes (miscellaneous) (no rank): 0.61Bacteroidetes bacterium (sp |
| 168 | 3.5 | 77.59 | 27.68 | 2 | Bacteria : 0.99FCB group (no rank): 0.67Bacteroidetes/Chlorobi group (no rank): 0.65Ignavibacteriae (phylum): 0.36Ignavibacteria (class): 0.31unclassified Ignavibacteria (no rank): 0.30 |
| 66 | 4.43 | 79.57 | 31.47 | 2 | Bacteria : 0.99unclassified Bacteria (no rank): 0.64Bacteria candidate phyla (no rank): 0.64Candidatus Rokubacteria (phylum): 0.62Candidatus Rokubacteria bacterium (species): 0.42 |
| 171 | 6 | 58.76 | 34.48 | 2 | Bacteria : 0.99FCB group (no rank): 0.66Gemmatimonadetes (phylum): 0.65unclassified Gemmatimonadetes (no rank): 0.35 |
| 58 | 4.61 | 78.88 | 38.45 | 2 | Bacteria : 0.99Nitrospirae (phylum): 0.81Nitrospira (class): 0.57Nitrospirales (order): 0.57Nitrospiraceae (family): 0.57Nitrospira (genus): 0.52 |
| 24 | 1.39 | 74.84 | 42.18 | 2 | Bacteria : 0.99unclassified Bacteria (no rank): 0.91Bacteria candidate phyla (no rank): 0.72Patescibacteria group (no rank): 0.71Parcubacteria group (no rank): 0.70Candidatus Moranbacteria (phylum): 0.64 |
| 53 | 1.82 | 79.15 | 44.97 | 2 | Bacteria : 0.98PVC group (no rank): 0.44Candidatus Omnitrophica (phylum): 0.43unclassified Candidatus Omnitrophica (no rank): 0.43 |
| 229 | 5.02 | 85.45 | 46.15 | 2 | Bacteria : 0.98unclassified Bacteria (no rank): 0.58Bacteria candidate phyla (no rank): 0.57Candidatus Eisenbacteria (phylum): 0.55 |
| 55 | 12.27 | 91.07 | 48.07 | 2 | Bacteria : 0.99Nitrospirae (phylum): 0.73Nitrospira (class): 0.34Nitrospirales (order): 0.34Nitrospiraceae (family): 0.34 |
| 166 | 5.44 | 85.58 | 58.79 | 2 | Bacteria : 0.99Terrabacteria group (no rank): 0.80Chloroflexi (phylum): 0.79 |
| 242 | 1.53 | 82.97 | 61.24 | 2 | Bacteria : 0.99unclassified Bacteria (no rank): 0.76Bacteria candidate phyla (no rank): 0.69Patescibacteria group (no rank): 0.67Parcubacteria group (no rank): 0.64Candidatus Moranbacteria (phylum): 0.48 |
| 264 | 5.22 | 82.19 | 63.17 | 2 | Bacteria : 0.99Proteobacteria (phylum): 0.83 |
| 176 | 3.12 | 83.01 | 63.83 | 2 | Archaea : 0.97TACK group (no rank): 0.97Thaumarchaeota (phylum): 0.95Nitrosopumilales (order): 0.79Nitrosopumilaceae (family): 0.36 |

| | | | | | | |
|---|---|---|---|---|---|---|
| 318 | 7.85 | 83.6 | 65.56 | 2 | | Bacteria : 0.99FCB group (no rank): 0.88Bacteroidetes/Chlorobi group (no rank): 0.87Bacteroidetes (phylum): 0.86unclassified Bacteroidetes (no rank): 0.65unclassified Bacteroidetes (miscellaneous) (no rank): 0.65 |
| 304 | 7.42 | 75.17 | 79.55 | 2 | | Bacteria : 0.99Proteobacteria (phylum): 0.81Betaproteobacteria (class): 0.63unclassified Betaproteobacteria (no rank): 0.51unclassified Betaproteobacteria (miscellaneous) (no rank): 0.51Betaproteobacteria bacterium (species): 0.35 |
| 235 | 1.99 | 72.35 | 79.99 | 2 | | Archaea : 0.80 |
| 125 | 6.92 | 94.91 | 81.05 | 2 | | Bacteria : 0.99FCB group (no rank): 0.59Gemmatimonadetes (phylum): 0.58 |
| 187 | 9.11 | 96.55 | 82.76 | 2 | | Bacteria : 0.97Proteobacteria (phylum): 0.75delta/epsilon subdivisions (subphylum): 0.73Deltaproteobacteria (class): 0.73unclassified Deltaproteobacteria (no rank): 0.61unclassified Deltaproteobacteria (miscellaneous) (no rank): 0.61Deltapr |
| 34 | 14.22 | 80.46 | 100.8 | 2 | | Bacteria : 0.99PVC group (no rank): 0.56Verrucomicrobia (phylum): 0.43 |
| 1 | 9.67 | 97.41 | 102.6 | 2 | | Bacteria : 0.99Proteobacteria (phylum): 0.68delta/epsilon subdivisions (subphylum): 0.61Deltaproteobacteria (class): 0.61unclassified Deltaproteobacteria (no rank): 0.51unclassified Deltaproteobacteria (miscellaneous) (no rank): 0.51Deltapr |
| 170 | 13 | 93.1 | 107.6 | 2 | | Bacteria : 0.99Proteobacteria (phylum): 0.73Betaproteobacteria (class): 0.33 |
| 268 | 9.68 | 93.1 | 108.2 | 2 | mmoC | Bacteria : 0.98Acidobacteria (phylum): 0.51unclassified Acidobacteria (no rank): 0.47 |
| 68 | 15.56 | 90.41 | 117 | 2 | pmoB | Bacteria : 0.99Proteobacteria (phylum): 0.83Betaproteobacteria (class): 0.61unclassified Betaproteobacteria (no rank): 0.51unclassified Betaproteobacteria (miscellaneous) (no rank): 0.50 |
| 69 | 13.07 | 96.39 | 190.1 | 2 | | Bacteria : 0.99Acidobacteria (phylum): 0.68unclassified Acidobacteria (no rank): 0.63unclassified Acidobacteria (miscellaneous) (no rank): 0.63 |
| 257 | 15.13 | 100 | 203.1 | 2 | | Bacteria : 0.99Proteobacteria (phylum): 0.85Alphaproteobacteria (class): 0.69Rhizobiales (order): 0.39 |
| 200 | 20.58 | 95.83 | 259.9 | 2 | | Bacteria : 0.99Proteobacteria (phylum): 0.78Betaproteobacteria (class): 0.57unclassified Betaproteobacteria (no rank): 0.44unclassified Betaproteobacteria (miscellaneous) (no rank): 0.44 |
| 227 | 15.94 | 98.28 | 294.7 | 2 | | Bacteria : 0.99Acidobacteria (phylum): 0.77unclassified Acidobacteria (no rank): 0.65unclassified Acidobacteria (miscellaneous) (no rank): 0.65Acidobacteria bacterium (species): 0.42 |
| 313 | 27.59 | 100 | 351.4 | 2 | mmoC | Bacteria : 0.99Acidobacteria (phylum): 0.68unclassified Acidobacteria (no rank): 0.61unclassified Acidobacteria (miscellaneous) (no rank): 0.61Acidobacteria bacterium (species): 0.35 |
| 2 | 29.97 | 100 | 352.2 | 2 | | Bacteria : 0.98Acidobacteria (phylum): 0.32 |
| 251 | 4.54 | 49.69 | 16.61 | 2 | | Bacteria : 0.99Acidobacteria (phylum): 0.53unclassified Acidobacteria (no rank): 0.42unclassified Acidobacteria (miscellaneous) (no rank): 0.42 |
| 329 | 2.57 | 49.32 | 8.72 | 2 | | Bacteria : 0.97Proteobacteria (phylum): 0.91Betaproteobacteria (class): 0.78Burkholderiales (order): 0.39 |
| 137 | 2.68 | 49.14 | 1.72 | 2 | | Bacteria : 0.99FCB group (no rank): 0.91Bacteroidetes/Chlorobi group (no rank): 0.91Bacteroidetes (phylum): 0.90unclassified Bacteroidetes (no rank): 0.69unclassified Bacteroidetes (miscellaneous) (no rank): 0.69 |
| 84 | 1.94 | 48.81 | 15.2 | 2 | | Bacteria : 1.00Terrabacteria group (no rank): 0.76Actinobacteria (phylum): 0.72Actinobacteria (class): 0.61unclassified Actinobacteria (class) (no rank): 0.54unclassified Actinobacteria (class) (miscellaneous) (no rank): 0.54Actinobacteria bacterium (species): 0.31 |
| 299 | 1.65 | 48.76 | 1.2 | 2 | | Archaea : 0.61Euryarchaeota (phylum): 0.46 |
| 129 | 3.41 | 48.47 | 0.91 | 2 | | Bacteria : 0.96Terrabacteria group (no rank): 0.64Chloroflexi (phylum): 0.60 |
| 301 | 1.83 | 48.28 | 5.17 | 2 | | Bacteria : 0.99FCB group (no rank): 0.57Gemmatimonadetes (phylum): 0.56 |
| 164 | 1.59 | 48.25 | 4.86 | 2 | | Bacteria : 0.99Terrabacteria group (no rank): 0.50Chloroflexi (phylum): 0.44unclassified Chloroflexi (no rank): 0.33unclassified Chloroflexi (miscellaneous) (no rank): 0.33 |
| 228 | 0.89 | 48.12 | 2.83 | 2 | | Bacteria : 0.98Terrabacteria group (no rank): 0.87Cyanobacteria/Melainabacteria group (no rank): 0.86Candidatus Saganbacteria (no rank): 0.86 |
| 293 | 1.16 | 47.89 | 0.91 | 2 | | Bacteria : 0.99Nitrospirae (phylum): 0.72 |
| 83 | 2.26 | 47.76 | 0 | 2 | | Bacteria : 0.99Proteobacteria (phylum): 0.87Gammaproteobacteria (class): 0.68unclassified Gammaproteobacteria (no rank): 0.56unclassified Gammaproteobacteria (miscellaneous) (no rank): 0.56Gammaproteobacteria bacterium (species): |
| 173 | 0.31 | 47.69 | 0.93 | 2 | | Archaea : 0.81 |
| 126 | 2.51 | 47.33 | 5.81 | 2 | | Bacteria : 0.99Proteobacteria (phylum): 0.64delta/epsilon subdivisions (subphylum): 0.45Deltaproteobacteria (class): 0.45unclassified Deltaproteobacteria (no rank): 0.40unclassified Deltaproteobacteria (miscellaneous) (no rank): 0.40Deltapr |
| 174 | 2.11 | 45.91 | 4.97 | 2 | mmoC | Bacteria : 0.99Nitrospirae (phylum): 0.59unclassified Nitrospirae (no rank): 0.34 |
| 71 | 0.36 | 45.65 | 0 | 2 | | Archaea : 0.75 |
| 192 | 0.86 | 45.57 | 0 | 2 | | Bacteria : 0.94 |
| 118 | 3.54 | 45.05 | 14.59 | 2 | | Bacteria : 0.99PVC group (no rank): 0.76Verrucomicrobia (phylum): 0.70unclassified Verrucomicrobia (no rank): 0.43unclassified Verrucomicrobia (miscellaneous) (no rank): 0.43Verrucomicrobia bacterium (species): 0.42 |
| 13 | 2.7 | 44.59 | 7.76 | 2 | | Bacteria : 0.99Terrabacteria group (no rank): 0.87Chloroflexi (phylum): 0.86 |
| 133 | 0.69 | 44.45 | 2.25 | 2 | | Bacteria : 0.98PVC group (no rank): 0.38Candidatus Omnitrophica (phylum): 0.38unclassified Candidatus Omnitrophica (no rank): 0.38 |
| 179 | 4.36 | 43.94 | 15.79 | 2 | mmoC | Bacteria : 0.99Nitrospirae (phylum): 0.83Nitrospira (class): 0.31Nitrospirales (order): 0.31Nitrospiraceae (family): 0.31 |
| 155 | 1.96 | 43.73 | 5.17 | 2 | | Bacteria : 0.99Acidobacteria (phylum): 0.53unclassified Acidobacteria (no rank): 0.49 |
| 321 | 1.98 | 42.46 | 2.58 | 2 | | Bacteria : 0.99Proteobacteria (phylum): 0.86delta/epsilon subdivisions (subphylum): 0.86Deltaproteobacteria (class): 0.86unclassified Deltaproteobacteria (no rank): 0.79unclassified Deltaproteobacteria (miscellaneous) (no rank): 0.79Deltapr |
| 263 | 0.65 | 42.15 | 1.98 | 2 | | Bacteria : 0.99unclassified Bacteria (no rank): 0.64Bacteria candidate phyla (no rank): 0.64Patescibacteria group (no rank): 0.63Microgenomates group (no rank): 0.60unclassified Microgenomates group (no rank): 0.38 |
| 148 | 2.61 | 42.02 | 0.07 | 2 | | Bacteria : 0.98Proteobacteria (phylum): 0.74delta/epsilon subdivisions (subphylum): 0.71Deltaproteobacteria (class): 0.71unclassified Deltaproteobacteria (no rank): 0.57unclassified Deltaproteobacteria (miscellaneous) (no rank): 0.57 |
| 7 | 1.17 | 41.22 | 5.06 | 2 | | Bacteria : 0.97Terrabacteria group (no rank): 0.87Chloroflexi (phylum): 0.86Dehalococcoidia (class): 0.42unclassified Dehalococcoidia (no rank): 0.41Dehalococcoidia bacterium (species): 0.39 |
| 146 | 1.37 | 40.93 | 0.55 | 2 | | Bacteria : 0.98FCB group (no rank): 0.53Bacteroidetes/Chlorobi group (no rank): 0.49 |
| 306 | 0.93 | 40.66 | 0.08 | 2 | | Bacteria : 1.00Nitrospirae (phylum): 0.66Nitrospira (class): 0.63Nitrospirales (order): 0.60Nitrospiraceae (family): 0.60unclassified Nitrospiraceae (no rank): 0.60Nitrospiraceae bacterium* (species): 0.60 |
| 208 | 0.61 | 40.6 | 0.85 | 2 | | Bacteria : 1.00unclassified Bacteria (no rank): 0.93Bacteria candidate phyla (no rank): 0.93candidate division NC10 (phylum): 0.91Candidatus Methylomirabilis (genus): 0.77Candidatus Methylomirabilis limnetica (species): 0.67 |
| 30 | 0.56 | 39.17 | 1.19 | 2 | | Bacteria : 0.99 |
| 70 | 2.5 | 39.17 | 2.55 | 2 | pmoA | Bacteria : 0.98Proteobacteria (phylum): 0.70 |
| 107 | 1.75 | 39.14 | 0 | 2 | | Bacteria : 1.00FCB group (no rank): 0.79Gemmatimonadetes (phylum): 0.78unclassified Gemmatimonadetes (no rank): 0.70Gemmatimonadetes bacterium (species): 0.66 |
| 260 | 0.67 | 39.14 | 18.84 | 2 | | Bacteria : 1.00unclassified Bacteria (no rank): 0.68Bacteria candidate phyla (no rank): 0.67Patescibacteria group (no rank): 0.64Microgenomates group (no rank): 0.59 |
| 233 | 1.34 | 38.66 | 0 | 2 | | Bacteria : 0.97PVC group (no rank): 0.70Planctomycetes (phylum): 0.60 |
| 311 | 1.45 | 38.45 | 2.75 | 2 | | Bacteria : 0.98Terrabacteria group (no rank): 0.73Chloroflexi (phylum): 0.69unclassified Chloroflexi (no rank): 0.41unclassified Chloroflexi (miscellaneous) (no rank): 0.41 |
| 158 | 0.95 | 37.65 | 0 | 2 | | Bacteria : 0.98unclassified Bacteria (no rank): 0.81Bacteria candidate phyla (no rank): 0.80Candidatus Edwardsbacteria (phylum): 0.77 |
| 217 | 0.32 | 37.16 | 0 | 2 | | Bacteria : 0.98unclassified Bacteria (no rank): 0.94Bacteria candidate phyla (no rank): 0.81Patescibacteria group (no rank): 0.81Parcubacteria group (no rank): 0.77Candidatus Jorgensenbacteria (phylum): 0.54 |
| 72 | 2.16 | 37.16 | 3.3 | 2 | | Bacteria : 1.00FCB group (no rank): 0.66Gemmatimonadetes (phylum): 0.65unclassified Gemmatimonadetes (no rank): 0.43Gemmatimonadetes bacterium (species): 0.31 |
| 336 | 1.07 | 36.95 | 2.84 | 2 | | Bacteria : 0.98PVC group (no rank): 0.68Planctomycetes (phylum): 0.67unclassified Planctomycetes (no rank): 0.35 |
| 76 | 0.39 | 36.88 | 4.52 | 2 | | Archaea : 0.89 |
| 12 | 1.03 | 36.55 | 5.96 | 2 | | Bacteria : 0.97PVC group (no rank): 0.51Verrucomicrobia (phylum): 0.40 |
| 128 | 1.14 | 36.28 | 1.91 | 2 | | Bacteria : 0.99FCB group (no rank): 0.36Bacteroidetes/Chlorobi group (no rank): 0.35 |
| 220 | 0.48 | 36.2 | 0.86 | 2 | | Bacteria : 0.97unclassified Bacteria (no rank): 0.93Bacteria candidate phyla (no rank): 0.91Patescibacteria group (no rank): 0.88Parcubacteria group (no rank): 0.84Candidatus Staskawiczbacteria (phylum): 0.35 |
| 89 | 0.44 | 35.72 | 0 | 2 | | Archaea : 0.96TACK group (no rank): 0.92Candidatus Bathyarchaeota (phylum): 0.91unclassified Candidatus Bathyarchaeota (no rank): 0.62Candidatus Bathyarchaeota archaeon (species): 0.49 |
| 266 | 1.21 | 35.62 | 2.07 | 2 | | Bacteria : 1.00PVC group (no rank): 0.81Verrucomicrobia (phylum): 0.79unclassified Verrucomicrobia (no rank): 0.46unclassified Verrucomicrobia (miscellaneous) (no rank): 0.46Verrucomicrobia bacterium (species): 0.46 |
| 215 | 1.27 | 35.02 | 0.55 | 2 | | Bacteria : 0.98FCB group (no rank): 0.59Bacteroidetes/Chlorobi group (no rank): 0.56 |

| | | | | | | |
|---|---|---|---|---|---|---|
| 186 | 0.64 | 34.86 | 0.03 | 2 | | Bacteria : 0.97Terrabacteria group (no rank): 0.89Firmicutes (phylum): 0.86unclassified Firmicutes sensu stricto (no rank): 0.74unclassified Firmicutes sensu stricto (miscellaneous) (no rank): 0.74Firmicutes bacterium (species): 0.74 |
| 285 | 1.46 | 34.09 | 7.76 | 2 | | Bacteria : 1.00Proteobacteria (phylum): 0.89Gammaproteobacteria (class): 0.60unclassified Gammaproteobacteria (no rank): 0.51unclassified Gammaproteobacteria (miscellaneous) (no rank): 0.51Gammaproteobacteria bacterium (species): |
| 142 | 0.56 | 33.87 | 3.74 | 2 | | Archaea : 0.91TACK group (no rank): 0.87Candidatus Bathyarchaeota (phylum): 0.85unclassified Candidatus Bathyarchaeota (no rank): 0.61Candidatus Bathyarchaeota archaeon (species): 0.49 |
| 163 | 1.72 | 33.65 | 0 | 2 | | Bacteria : 0.98Proteobacteria (phylum): 0.78Gammaproteobacteria (class): 0.63Xanthomonadales (order): 0.34unclassified Xanthomonadales (no rank): 0.33Xanthomonadales bacterium (species): 0.33 |
| 262 | 1.14 | 33.21 | 0 | 2 | | Bacteria : 0.92 |
| 28 | 1.26 | 33.15 | 2.35 | 2 | mmoC | Bacteria : 0.87unclassified Bacteria (no rank): 0.51Bacteria candidate phyla (no rank): 0.49 |
| 138 | 1.69 | 32.76 | 0 | 2 | | Bacteria : 0.99Proteobacteria (phylum): 0.84 |
| 139 | 1.35 | 32.46 | 2.91 | 2 | | Bacteria : 0.95Proteobacteria (phylum): 0.64delta/epsilon subdivisions (subphylum): 0.63Deltaproteobacteria (class): 0.63 |
| 234 | 0.94 | 32.13 | 0 | 2 | | Bacteria : 0.99FCB group (no rank): 0.85candidate division Zixibacteria (phylum): 0.83candidate division Zixibacteria bacterium RBG_16_53_22 (species): 0.63 |
| 216 | 0.75 | 32.08 | 1.14 | 2 | | Bacteria : 0.97PVC group (no rank): 0.70Planctomycetes (phylum): 0.68unclassified Planctomycetes (no rank): 0.31 |
| 180 | 1.21 | 31.73 | 0.97 | 2 | | Bacteria : 0.99Proteobacteria (phylum): 0.90delta/epsilon subdivisions (subphylum): 0.89Deltaproteobacteria (class): 0.89Desulfobacterales (order): 0.79Desulfobacteraceae (family): 0.78unclassified Desulfobacteraceae (no rank): 0.72Desulf |
| 278 | 0.47 | 31.66 | 0.94 | 2 | | Bacteria : 0.96Proteobacteria (phylum): 0.62delta/epsilon subdivisions (subphylum): 0.62Deltaproteobacteria (class): 0.62 |
| 15 | 0.7 | 31.57 | 0 | 2 | | Bacteria : 1.00Acidobacteria (phylum): 0.87unclassified Acidobacteria (no rank): 0.85unclassified Acidobacteria (miscellaneous) (no rank): 0.85Acidobacteria bacterium RBG_13_68_16 (species): 0.64 |
| 105 | 0.97 | 31.03 | 1.72 | 2 | | Bacteria : 0.98Proteobacteria (phylum): 0.69Alphaproteobacteria (class): 0.52 |
| 291 | 0.23 | 30.41 | 0.57 | 2 | | Bacteria : 0.97unclassified Bacteria (no rank): 0.76Bacteria candidate phyla (no rank): 0.69Patescibacteria group (no rank): 0.64Parcubacteria group (no rank): 0.57 |
| 193 | 0.71 | 30.28 | 10.75 | 2 | | Archaea : 0.84environmental samples (no rank): 0.32uncultured archaeon (species): 0.32 |
| 283 | 1.57 | 30.24 | 0 | 2 | | Bacteria : 0.98Spirochaetes (phylum): 0.68unclassified Spirochaetes (no rank): 0.63Spirochaetes bacterium RBG_16_67_19 (species): 0.31 |
| 198 | 1.38 | 29.96 | 3.36 | 2 | | Bacteria : 1.00FCB group (no rank): 0.68Gemmatimonadetes (phylum): 0.67unclassified Gemmatimonadetes (no rank): 0.66 |
| 112 | 1.85 | 29.7 | 0 | 2 | | Bacteria : 0.99unclassified Bacteria (no rank): 0.65Bacteria candidate phyla (no rank): 0.65candidate division NC10 (phylum): 0.59unclassified candidate division NC10 (no rank): 0.59 |
| 35 | 1.47 | 29.64 | 0 | 2 | | Bacteria : 0.96Terrabacteria group (no rank): 0.66Chloroflexi (phylum): 0.62 |
| 305 | 1 | 29.19 | 4.22 | 2 | | Bacteria : 1.00Proteobacteria (phylum): 0.96delta/epsilon subdivisions (subphylum): 0.95Deltaproteobacteria (class): 0.95unclassified Deltaproteobacteria (no rank): 0.47unclassified Deltaproteobacteria (miscellaneous) (no rank): 0.47 |
| 290 | 1.1 | 29.08 | 1.99 | 2 | | Bacteria : 0.98unclassified Bacteria (no rank): 0.36Bacteria candidate phyla (no rank): 0.36candidate division NC10 (phylum): 0.31unclassified candidate division NC10 (no rank): 0.30 |
| 280 | 2.01 | 28.68 | 0 | 2 | | Bacteria : 0.97FCB group (no rank): 0.31 |
| 99 | 0.69 | 28.62 | 0 | 2 | | Bacteria : 0.99Terrabacteria group (no rank): 0.73Actinobacteria (phylum): 0.65 |
| 226 | 1.03 | 28.61 | 0.91 | 2 | | Bacteria : 0.99Terrabacteria group (no rank): 0.89Chloroflexi (phylum): 0.88Anaerolineae (class): 0.49 |
| 199 | 1.91 | 28.45 | 3.45 | 2 | | Bacteria : 0.99FCB group (no rank): 0.78Bacteroidetes/Chlorobi group (no rank): 0.77Bacteroidetes (phylum): 0.77 |
| 159 | 1.97 | 28.34 | 2.73 | 2 | | Bacteria : 0.98Acidobacteria (phylum): 0.50unclassified Acidobacteria (no rank): 0.45unclassified Acidobacteria (miscellaneous) (no rank): 0.45Acidobacteria bacterium (species): 0.41 |
| 110 | 1.03 | 27.83 | 0.54 | 2 | | Bacteria : 0.98FCB group (no rank): 0.80Bacteroidetes/Chlorobi group (no rank): 0.77Bacteroidetes (phylum): 0.75unclassified Bacteroidetes (no rank): 0.35unclassified Bacteroidetes (miscellaneous) (no rank): 0.35 |
| 230 | 0.9 | 26.55 | 0.13 | 2 | | Bacteria : 0.97unclassified Bacteria (no rank): 0.58Bacteria candidate phyla (no rank): 0.47candidate division KSB1 (no rank): 0.45unclassified candidate division KSB1 (no rank): 0.45candidate division KSB1 bacterium (species): 0.45 |
| 275 | 1.13 | 25.61 | 3.45 | 2 | | Bacteria : 1.00Proteobacteria (phylum): 0.92Gammaproteobacteria (class): 0.72unclassified Gammaproteobacteria (no rank): 0.62unclassified Gammaproteobacteria (miscellaneous) (no rank): 0.62Gammaproteobacteria bacterium (species): |
| 271 | 2.07 | 25.08 | 3.45 | 2 | | Bacteria : 1.00Proteobacteria (phylum): 0.90Betaproteobacteria (class): 0.76unclassified Betaproteobacteria (no rank): 0.67unclassified Betaproteobacteria (miscellaneous) (no rank): 0.67Betaproteobacteria bacterium RBG_16_66_20 (specie |
| 16 | 0.85 | 24.41 | 0 | 2 | | Bacteria : 0.96Terrabacteria group (no rank): 0.65Chloroflexi (phylum): 0.63unclassified Chloroflexi (no rank): 0.39unclassified Chloroflexi (miscellaneous) (no rank): 0.38 |
| 222 | 0.94 | 24.06 | 0 | 2 | | Bacteria : 0.99Proteobacteria (phylum): 0.84delta/epsilon subdivisions (subphylum): 0.81Deltaproteobacteria (class): 0.81Syntrophobacterales (order): 0.42 |
| 122 | 1.38 | 23.68 | 0 | 2 | | Bacteria : 1.00Proteobacteria (phylum): 0.68delta/epsilon subdivisions (subphylum): 0.63Deltaproteobacteria (class): 0.63unclassified Deltaproteobacteria (no rank): 0.58unclassified Deltaproteobacteria (miscellaneous) (no rank): 0.58Deltapr |
| 324 | 0.24 | 23.43 | 0 | 2 | | Bacteria : 1.00Proteobacteria (phylum): 0.95Gammaproteobacteria (class): 0.76unclassified Gammaproteobacteria (no rank): 0.71unclassified Gammaproteobacteria (miscellaneous) (no rank): 0.71Gammaproteobacteria bacterium (species): |
| 123 | 0.42 | 22.72 | 0 | 2 | | Archaea : 0.80DPANN group (no rank): 0.52Candidatus Woesearchaeota (phylum): 0.45 |
| 152 | 0.37 | 22.41 | 0 | 2 | | Bacteria : 0.99Proteobacteria (phylum): 0.74Betaproteobacteria (class): 0.32 |
| 17 | 1.44 | 21.47 | 0 | 2 | | Bacteria : 0.98FCB group (no rank): 0.58Bacteroidetes/Chlorobi group (no rank): 0.56 |
| 240 | 0.31 | 21.22 | 0 | 2 | | Archaea : 0.91environmental samples (no rank): 0.85uncultured archaeon (species): 0.85 |
| 38 | 1.36 | 21.22 | 0.16 | 2 | | Bacteria : 0.99Terrabacteria group (no rank): 0.89Chloroflexi (phylum): 0.88 |
| 255 | 0.78 | 21.05 | 0 | 2 | | Bacteria : 1.00FCB group (no rank): 0.63Gemmatimonadetes (phylum): 0.62unclassified Gemmatimonadetes (no rank): 0.39 |
| 45 | 0.36 | 21.04 | 0 | 2 | | Archaea : 0.98TACK group (no rank): 0.98Thaumarchaeota (phylum): 0.97Nitrosopumilales (order): 0.94Nitrosopumilaceae (family): 0.67 |
| 205 | 1.15 | 20.77 | 1.18 | 2 | | Bacteria : 0.99Terrabacteria group (no rank): 0.88Chloroflexi (phylum): 0.87Anaerolineae (class): 0.45 |
| 108 | 0.65 | 20.58 | 0 | 2 | | Bacteria : 0.98PVC group (no rank): 0.67Planctomycetes (phylum): 0.67 |
| 279 | 0.41 | 20.33 | 0.47 | 2 | | Archaea : 0.88environmental samples (no rank): 0.63uncultured archaeon (species): 0.63 |
| 177 | 1.14 | 20.06 | 0 | 2 | | Bacteria : 0.99unclassified Bacteria (no rank): 0.88Bacteria candidate phyla (no rank): 0.84Candidatus Aminicenantes (phylum): 0.84unclassified Aminicenantes (no rank): 0.83 |
| 287 | 0.47 | 19.83 | 0 | 2 | | Bacteria : 1.00Proteobacteria (phylum): 0.95delta/epsilon subdivisions (subphylum): 0.94Deltaproteobacteria (class): 0.94unclassified Deltaproteobacteria (no rank): 0.93unclassified Deltaproteobacteria (miscellaneous) (no rank): 0.93 |
| 136 | 0.69 | 19.03 | 0 | 2 | | Bacteria : 0.99FCB group (no rank): 0.70Gemmatimonadetes (phylum): 0.69 |
| 62 | 0.28 | 18.96 | 0.81 | 2 | | Archaea : 0.89 |
| 273 | 0.85 | 18.92 | 0 | 2 | | Bacteria : 1.00Proteobacteria (phylum): 0.91Betaproteobacteria (class): 0.81unclassified Betaproteobacteria (no rank): 0.78unclassified Betaproteobacteria (miscellaneous) (no rank): 0.78Betaproteobacteria bacterium RBG_16_66_20 (specie |
| 132 | 2.2 | 18.79 | 6.9 | 2 | | Bacteria : 0.99unclassified Bacteria (no rank): 0.73Bacteria candidate phyla (no rank): 0.72Candidatus Rokubacteria (phylum): 0.70Candidatus Rokubacteria bacterium (species): 0.46 |
| 212 | 0.62 | 18.79 | 1.82 | 2 | | Bacteria : 0.98Terrabacteria group (no rank): 0.83Chloroflexi (phylum): 0.81 |
| 247 | 0.45 | 17.24 | 3.45 | 2 | | Bacteria : 0.98Proteobacteria (phylum): 0.83 |
| 331 | 0.68 | 17.19 | 0.65 | 2 | | Bacteria : 0.98Proteobacteria (phylum): 0.72delta/epsilon subdivisions (subphylum): 0.69Deltaproteobacteria (class): 0.69 |
| 65 | 0.29 | 17.09 | 0 | 2 | | Bacteria : 1.00PVC group (no rank): 0.95Planctomycetes (phylum): 0.94unclassified Planctomycetes (no rank): 0.93Planctomycetes bacterium GWC2_45_44 (species): 0.63 |
| 52 | 0.46 | 17.01 | 1.72 | 2 | | Bacteria : 1.00Terrabacteria group (no rank): 0.90Chloroflexi (phylum): 0.89unclassified Chloroflexi (no rank): 0.39unclassified Chloroflexi (miscellaneous) (no rank): 0.39 |
| 61 | 0.34 | 16.83 | 0 | 2 | | Archaea : 0.92TACK group (no rank): 0.89Thaumarchaeota (phylum): 0.87Nitrosopumilales (order): 0.62 |
| 337 | 0.82 | 15.86 | 0 | 2 | | Bacteria : 0.98Proteobacteria (phylum): 0.71Alphaproteobacteria (class): 0.59Rhodospirillales (order): 0.35Rhodospirillaceae (family): 0.31 |
| 106 | 0.46 | 15.79 | 0 | 2 | | Bacteria : 1.00Nitrospirae (phylum): 0.93Nitrospira (class): 0.42Nitrospirales (order): 0.42Nitrospiraceae (family): 0.42 |
| 115 | 0.39 | 15.58 | 0 | 2 | | Archaea : 0.60DPANN group (no rank): 0.32 |
| 269 | 0.38 | 15.27 | 0 | 2 | | Bacteria : 0.90 |

| | | | | | |
|---|---|---|---|---|---|
| 77 | 0.43 | 14.89 | 0 | 2 | Bacteria : 0.98FCB group (no rank): 0.63Candidatus Latescibacteria (phylum): 0.58unclassified Candidatus Latescibacteria (no rank): 0.58Candidatus Latescibacteria bacterium (species): 0.49 |
| 327 | 0.4 | 14.35 | 1.75 | 2 | Bacteria : 0.95Terrabacteria group (no rank): 0.89Chloroflexi (phylum): 0.88Dehalococcoidia (class): 0.42unclassified Dehalococcoidia (no rank): 0.41Dehalococcoidia bacterium (species): 0.40 |
| 243 | 0.47 | 12.78 | 0 | 2 | Bacteria : 0.98PVC group (no rank): 0.42Planctomycetes (phylum): 0.33 |
| 31 | 0.59 | 12.5 | 0 | 2 | Bacteria : 1.00FCB group (no rank): 0.64Gemmatimonadetes (phylum): 0.64 |
| 63 | 0.21 | 12.15 | 0 | 2 | Archaea : 0.85Euryarchaeota (phylum): 0.60unclassified Euryarchaeota (no rank): 0.56unclassified Euryarchaeota (miscellaneous) (no rank): 0.56Euryarchaeota archaeon SM23-78 (species): 0.56 |
| 9 | 0.51 | 12.07 | 0 | 2 | Bacteria : 0.96Proteobacteria (phylum): 0.89Betaproteobacteria (class): 0.79Burkholderiales (order): 0.32 |
| 64 | 0.33 | 11.69 | 0 | 2 | Bacteria : 0.98Terrabacteria group (no rank): 0.80Chloroflexi (phylum): 0.77 |
| 308 | 0.3 | 11.13 | 0 | 2 | Bacteria : 1.00Proteobacteria (phylum): 0.84 |
| 258 | 0.76 | 10.85 | 0 | 2 | Bacteria : 0.98Nitrospirae (phylum): 0.81Nitrospira (class): 0.34Nitrospirales (order): 0.34Nitrospiraceae (family): 0.34 |
| 82 | 6.28 | 10.75 | 16.02 | 2 | Bacteria : 0.46 |
| 207 | 1.12 | 10.68 | 1.94 | 2 | Archaea : 0.82TACK group (no rank): 0.74Candidatus Bathyarchaeota (phylum): 0.72unclassified Candidatus Bathyarchaeota (no rank): 0.65Candidatus Bathyarchaeota archaeon (species): 0.57 |
| 120 | 0.92 | 10.42 | 0 | 2 | Bacteria : 0.99Proteobacteria (phylum): 0.71Gammaproteobacteria (class): 0.49unclassified Gammaproteobacteria (no rank): 0.41unclassified Gammaproteobacteria (miscellaneous) (no rank): 0.35Gammaproteobacteria bacterium (species): |
| 19 | 0.6 | 10.03 | 0.94 | 2 | Bacteria : 1.00Proteobacteria (phylum): 0.90Betaproteobacteria (class): 0.84unclassified Betaproteobacteria (no rank): 0.81unclassified Betaproteobacteria (miscellaneous) (no rank): 0.81 |
| 325 | 0.42 | 9.84 | 0 | 2 | Bacteria : 1.00Proteobacteria (phylum): 0.86Betaproteobacteria (class): 0.37 |
| 80 | 1.6 | 9.53 | 1.38 | 2 | Bacteria : 0.98PVC group (no rank): 0.50Verrucomicrobia (phylum): 0.44 |
| 254 | 0.23 | 8.78 | 0 | 2 | Bacteria : 0.99Acidobacteria (phylum): 0.54unclassified Acidobacteria (no rank): 0.54unclassified Acidobacteria (miscellaneous) (no rank): 0.51Acidobacteria bacterium (species): 0.42 |
| 312 | 1.32 | 8.33 | 4.17 | 2 | Bacteria : 0.99Proteobacteria (phylum): 0.81 |
| 102 | 0.27 | 7.94 | 0 | 2 | Bacteria : 0.86Terrabacteria group (no rank): 0.65Chloroflexi (phylum): 0.61 |
| 326 | 0.33 | 7.52 | 0 | 2 | Bacteria : 0.98Proteobacteria (phylum): 0.92Betaproteobacteria (class): 0.76Nitrosomonadales (order): 0.31 |
| 100 | 0.34 | 6.97 | 0 | 2 | Bacteria : 0.87unclassified Bacteria (no rank): 0.55Bacteria candidate phyla (no rank): 0.54 |
| 288 | 0.5 | 6.9 | 0 | 2 | Bacteria : 0.99FCB group (no rank): 0.64Gemmatimonadetes (phylum): 0.63unclassified Gemmatimonadetes (no rank): 0.52 |
| 49 | 0.3 | 6.58 | 0 | 2 | Bacteria : 0.99Acidobacteria (phylum): 0.64 |
| 211 | 0.4 | 6.43 | 0 | 2 | Bacteria : 1.00Terrabacteria group (no rank): 0.88Chloroflexi (phylum): 0.79unclassified Chloroflexi (no rank): 0.49unclassified Chloroflexi (miscellaneous) (no rank): 0.49 |
| 94 | 0.91 | 6.35 | 0 | 2 | Bacteria : 0.99Terrabacteria group (no rank): 0.89Chloroflexi (phylum): 0.88 |
| 130 | 0.25 | 5.17 | 0 | 2 | Bacteria : 1.00Proteobacteria (phylum): 0.90Betaproteobacteria (class): 0.73unclassified Betaproteobacteria (no rank): 0.63unclassified Betaproteobacteria (miscellaneous) (no rank): 0.62 |
| 219 | 0.26 | 4.17 | 0 | 2 | Bacteria : 0.81 |
| 284 | 0.42 | 4.17 | 0 | 2 | Bacteria : 0.99FCB group (no rank): 0.53Bacteroidetes/Chlorobi group (no rank): 0.49 |
| 307 | 0.21 | 4.17 | 0 | 2 | Bacteria : 0.98Proteobacteria (phylum): 0.79delta/epsilon subdivisions (subphylum): 0.72Deltaproteobacteria (class): 0.72unclassified Deltaproteobacteria (no rank): 0.68unclassified Deltaproteobacteria (miscellaneous) (no rank): 0.68 |
| 42 | 0.24 | 4.17 | 0 | 2 | Bacteria : 1.00PVC group (no rank): 0.85Candidatus Omnitrophica (phylum): 0.83unclassified Candidatus Omnitrophica (no rank): 0.83Omnitrophica bacterium RIFOXYB12_FULL_50_7 (species): 0.63 |
| 48 | 0.46 | 4.17 | 0 | 2 | Bacteria : 1.00FCB group (no rank): 0.65Gemmatimonadetes (phylum): 0.64unclassified Gemmatimonadetes (no rank): 0.35 |
| 56 | 1.13 | 4.17 | 0 | 2 | Bacteria : 0.97Terrabacteria group (no rank): 0.77Chloroflexi (phylum): 0.75unclassified Chloroflexi (no rank): 0.59unclassified Chloroflexi (miscellaneous) (no rank): 0.59 |
| 44 | 0.49 | 4.09 | 0 | 2 | Bacteria : 0.94Proteobacteria (phylum): 0.78delta/epsilon subdivisions (subphylum): 0.76Deltaproteobacteria (class): 0.76Desulfobacterales (order): 0.65Desulfobacteraceae (family): 0.64unclassified Desulfobacteraceae (no rank): 0.62Desulf[illegible] |
| 259 | 0.64 | 3.7 | 0 | 2 | Bacteria : 0.41 |
| 79 | 0.37 | 3.62 | 0 | 2 | Bacteria : 0.52 |
| 332 | 0.32 | 3.06 | 1.15 | 2 | |
| 238 | 0.37 | 2.9 | 0 | 2 | Bacteria : 0.94PVC group (no rank): 0.55Verrucomicrobia (phylum): 0.53 |
| 214 | 0.63 | 2.68 | 0.38 | 2 | |
| 197 | 0.22 | 2.63 | 0 | 2 | Bacteria : 0.87 |
| 88 | 0.36 | 2.58 | 0.22 | 2 | |
| 36 | 0.29 | 2.43 | 0 | 2 | Bacteria : 0.93FCB group (no rank): 0.85Bacteroidetes/Chlorobi group (no rank): 0.85Ignavibacteriae (phylum): 0.79Ignavibacteria (class): 0.73Ignavibacteriales (order): 0.70unclassified Ignavibacteriales (no rank): 0.42 |
| 172 | 0.49 | 2.31 | 6.94 | 2 | Bacteria : 0.63 |
| 183 | 0.27 | 2.31 | 1.85 | 2 | Bacteria : 0.50 |
| 51 | 0.31 | 2.23 | 0 | 2 | Bacteria : 0.99Acidobacteria (phylum): 0.36unclassified Acidobacteria (no rank): 0.32unclassified Acidobacteria (miscellaneous) (no rank): 0.31 |
| 300 | 0.31 | 1.89 | 0 | 2 | Bacteria : 0.37 |
| 225 | 0.74 | 1.88 | 0.78 | 2 | Bacteria : 0.96Proteobacteria (phylum): 0.59delta/epsilon subdivisions (subphylum): 0.56Deltaproteobacteria (class): 0.56unclassified Deltaproteobacteria (no rank): 0.37unclassified Deltaproteobacteria (miscellaneous) (no rank): 0.37 |
| 224 | 0.27 | 1.21 | 0 | 2 | |
| 109 | 0.29 | 0.97 | 1.14 | 2 | Bacteria : 0.57 |
| 135 | 1.86 | 0.86 | 0 | 2 | Bacteria : 0.97Proteobacteria (phylum): 0.45delta/epsilon subdivisions (subphylum): 0.35Deltaproteobacteria (class): 0.35unclassified Deltaproteobacteria (no rank): 0.31unclassified Deltaproteobacteria (miscellaneous) (no rank): 0.31 |
| 113 | 0.3 | 0.78 | 0.47 | 2 | Bacteria : 0.61 |
| 292 | 0.25 | 0.72 | 0 | 2 | unclassified viruses : 0.71Pacmanvirus A23 (species): 0.61 |
| 141 | 0.23 | 0.53 | 0 | 2 | |
| 239 | 0.38 | 0.47 | 0 | 2 | Bacteria : 0.67 |
| 236 | 0.35 | 0.14 | 0 | 2 | Bacteria : 0.53 |
| 54 | 0.25 | 0.07 | 0 | 2 | Bacteria : 0.84 |
| 10 | 0.28 | 0 | 0 | 2 | Archaea : 0.87TACK group (no rank): 0.82Candidatus Bathyarchaeota (phylum): 0.79unclassified Candidatus Bathyarchaeota (no rank): 0.69Candidatus Bathyarchaeota archaeon (species): 0.65 |
| 104 | 0.42 | 0 | 0 | 2 | Bacteria : 0.50 |
| 11 | 1.82 | 0 | 0 | 2 | Bacteria : 0.98FCB group (no rank): 0.48Gemmatimonadetes (phylum): 0.46unclassified Gemmatimonadetes (no rank): 0.38 |
| 131 | 0.35 | 0 | 0 | 2 | Bacteria : 0.99Proteobacteria (phylum): 0.67delta/epsilon subdivisions (subphylum): 0.63Deltaproteobacteria (class): 0.63 |
| 144 | 0.32 | 0 | 0 | 2 | Bacteria : 0.53 |
| 150 | 0.42 | 0 | 0 | 2 | Bacteria : 0.96Nitrospirae (phylum): 0.33 |
| 151 | 0.25 | 0 | 0 | 2 | Bacteria : 0.69 |

| | | | | | |
|---|---|---|---|---|---|
| 161 | 0.3 | 0 | 0 | 2 | Bacteria : 0.36 |
| 162 | 0.23 | 0 | 0 | 2 | Bacteria : 0.71 |
| 189 | 0.28 | 0 | 0 | 2 | Bacteria : 0.96Proteobacteria (phylum): 0.56delta/epsilon subdivisions (subphylum): 0.50Deltaproteobacteria (class): 0.50 |
| 190 | 0.22 | 0 | 0 | 2 | Bacteria : 0.70 |
| 204 | 0.75 | 0 | 0 | 2 | Bacteria : 0.99Proteobacteria (phylum): 0.82 |
| 206 | 0.2 | 0 | 0 | 2 | Bacteria : 1.00Proteobacteria (phylum): 0.56 |
| 213 | 0.46 | 0 | 0 | 2 | Bacteria : 0.94Proteobacteria (phylum): 0.48delta/epsilon subdivisions (subphylum): 0.35Deltaproteobacteria (class): 0.35 |
| 218 | 0.37 | 0 | 0 | 2 | Bacteria : 0.96Proteobacteria (phylum): 0.38 |
| 22 | 0.31 | 0 | 0 | 2 | Bacteria : 0.40 |
| 221 | 0.47 | 0 | 0 | 2 | Bacteria : 0.94FCB group (no rank): 0.72Bacteroidetes/Chlorobi group (no rank): 0.72Bacteroidetes (phylum): 0.65 |
| 223 | 0.41 | 0 | 0 | 2 | Bacteria : 0.86 |
| 23 | 0.26 | 0 | 0 | 2 | Bacteria : 0.56 |
| 232 | 0.23 | 0 | 0 | 2 | Bacteria : 0.77 |
| 246 | 0.86 | 0 | 0 | 2 | Bacteria : 0.92Proteobacteria (phylum): 0.44delta/epsilon subdivisions (subphylum): 0.37Deltaproteobacteria (class): 0.37 |
| 248 | 0.28 | 0 | 0 | 2 | Bacteria : 0.59 |
| 250 | 0.28 | 0 | 0 | 2 | Bacteria : 0.33 |
| 252 | 0.39 | 0 | 0 | 2 | Bacteria : 0.31 |
| 261 | 0.21 | 0 | 0 | 2 | Bacteria : 0.94Terrabacteria group (no rank): 0.70Chloroflexi (phylum): 0.67 |
| 267 | 0.36 | 0 | 0 | 2 | Bacteria : 0.95Proteobacteria (phylum): 0.46delta/epsilon subdivisions (subphylum): 0.45Deltaproteobacteria (class): 0.45 |
| 27 | 0.24 | 0 | 0 | 2 | Bacteria : 0.57 |
| 272 | 0.35 | 0 | 0 | 2 | Bacteria : 0.84 |
| 29 | 0.23 | 0 | 0 | 2 | Eukaryota : 0.37 |
| 294 | 0.25 | 0 | 0 | 2 | Bacteria : 0.98Proteobacteria (phylum): 0.76delta/epsilon subdivisions (subphylum): 0.68Deltaproteobacteria (class): 0.68unclassified Deltaproteobacteria (no rank): 0.57unclassified Deltaproteobacteria (miscellaneous) (no rank): 0.57 |
| 302 | 0.58 | 0 | 0 | 2 | Bacteria : 0.84unclassified Bacteria (no rank): 0.35Bacteria candidate phyla (no rank): 0.34Patescibacteria group (no rank): 0.34 |
| 303 | 0.22 | 0 | 0 | 2 | Bacteria : 0.61 |
| 316 | 0.3 | 0 | 0 | 2 | Bacteria : 0.95Spirochaetes (phylum): 0.65unclassified Spirochaetes (no rank): 0.65Spirochaetes bacterium (species): 0.65 |
| 320 | 0.3 | 0 | 0 | 2 | Bacteria : 0.99unclassified Bacteria (no rank): 0.68Bacteria candidate phyla (no rank): 0.68Candidatus Rokubacteria (phylum): 0.66 |
| 43 | 0.33 | 0 | 0 | 2 | Bacteria : 0.60 |
| 57 | 0.24 | 0 | 0 | 2 | Bacteria : 0.95 |
| 59 | 0.6 | 0 | 0 | 2 | Bacteria : 0.93Proteobacteria (phylum): 0.57delta/epsilon subdivisions (subphylum): 0.54Deltaproteobacteria (class): 0.54unclassified Deltaproteobacteria (no rank): 0.38unclassified Deltaproteobacteria (miscellaneous) (no rank): 0.38 |
| 6 | 0.33 | 0 | 0 | 2 | Bacteria : 0.94Proteobacteria (phylum): 0.37 |
| 60 | 0.36 | 0 | 0 | 2 | Bacteria : 0.72Proteobacteria (phylum): 0.34 |
| 73 | 0.25 | 0 | 0 | 2 | Bacteria : 0.96unclassified Bacteria (no rank): 0.63Bacteria candidate phyla (no rank): 0.62Candidatus Eisenbacteria (phylum): 0.59 |
| 74 | 0.25 | 0 | 0 | 2 | |
| 81 | 0.26 | 0 | 0 | 2 | Bacteria : 0.80 |
| 87 | 0.28 | 0 | 0 | 2 | Bacteria : 0.99Proteobacteria (phylum): 0.58 |
| 93 | 0.34 | 0 | 0 | 2 | Bacteria : 0.98FCB group (no rank): 0.60Bacteroidetes/Chlorobi group (no rank): 0.58Ignavibacteriae (phylum): 0.31 |
